# SexFindR: A computational workflow to identify young and old sex chromosomes

**DOI:** 10.1101/2022.02.21.481346

**Authors:** Phil Grayson, Alison Wright, Colin J. Garroway, Margaret F. Docker

## Abstract

Sex chromosomes have evolved frequently across the tree of life, and have been a source of fascination for decades due to their unique evolutionary trajectories. They are hypothesised to be important drivers in a broad spectrum of biological processes and are the focus of a rich body of evolutionary theory. Whole-genome sequencing provides exciting opportunities to test these theories through contrasts between independently evolved sex chromosomes across the full spectrum of their evolutionary lifecycles. However, identifying sex chromosomes, particularly nascent ones, is challenging, often requiring specific combinations of methodologies. This is a major barrier to progress in the field and can result in discrepancies between studies that apply different approaches. Currently, no single pipeline exists to integrate data across these methods in a statistical framework to identify sex chromosomes at all ages and levels of sequence divergence. To address this, we present SexFindR, a comprehensive workflow to improve robustness and transparency in identifying sex-linked sequences. We validate our approach using publicly available data from five species that span the continuum of sex chromosome divergence, from homomorphic sex chromosomes with only a single SNP that determines sex, to heteromorphic sex chromosomes with extensive degeneration. Next, we apply SexFindR to our large-scale population genomics dataset for sea lamprey, a jawless vertebrate whose sex determination system remains a mystery despite decades of research. We decisively show that sea lamprey do not harbour sex-linked sequences in their somatic genome, leaving open the possibility that sex is determined environmentally or within the germline genome.

## Introduction

### Sex Chromosome Evolution

Beginning as an identical pair of autosomes, sex chromosomes evolve through the acquisition of a master sex-determining locus (Furman et al. 2020). Over time, recombination suppression can occur between the sex chromosome pair and, in many cases, will proceed along the chromosome length, resulting in highly divergent, heteromorphic sex chromosomes (Charlesworth et al. 2005; Wright et al. 2016; Vicoso 2019; Furman et al. 2020), where the heterogametic sex is the sex that carries two different sex chromosomes. However, sex chromosome divergence and degeneration are not inevitable, and numerous species of animals and plants exhibit homomorphic sex chromosomes that are largely identical in size and gene content (Vicoso 2019). Studying sex chromosomes at the early stages of divergence can provide valuable data, making it possible to explore the processes driving the formation of new sex chromosomes.

Despite a rich body of theoretical expectations (Palmer et al. 2019; Vicoso 2019; Furman et al. 2020), we are still far from a comprehensive understanding of the forces that catalyse sex chromosome formation and turnover. To robustly test these theories, we require contrasts between independently evolved sex chromosomes across the full spectrum of their evolutionary lifecycle. Recent advances in DNA sequencing and genome assembly, alongside reduction in overall costs has made this increasingly possible through data from numerous non-model organisms. Even so, identifying sex chromosomes from sequence data poses a number of challenges, particularly for nascent sex chromosomes at their early stages of divergence (Palmer et al. 2019). A number of pipelines do exist to identify sex chromosomes from sequence data, including SEX-DETector (Muyle et al. 2016), FindZX (Sigeman, Sinclair, et al. 2021), discoverY (Rangavittal et al. 2019), RADSex (Feron et al. 2021) and detsex (Gautier 2014). These pipelines employ a range of different methodologies, contrasting patterns of coverage (e.g., FindZX) or heterozygosity (e.g., FindZX, RADSex) between the sexes or segregation of alleles across a pedigree (e.g., SEX-DETector). These are powerful approaches to identify sex-linked sequences, however, they either require specific types of data, such as pedigree information, or *a priori* knowledge of the sex chromosome system, or are only able to detect sex chromosomes at certain stages of divergence. This is because as sex chromosomes diverge, they leave distinct signatures in sequence data which change as divergence proceeds, and so a combination of different methods is necessary to identify sex chromosomes among species (Palmer et al. 2019). However, we currently lack a statistical framework to robustly integrate the results of these different methods to identify a high confidence set of sex-linked sequences and determine if a species exhibits genetic sex determination across the full divergence gradient. This is a major barrier to progress as the use of different approaches can lead to discrepancies between studies as a result of methodological biases (Darolti et al. 2021).

### Current Approaches for Sex Chromosome Identification

Heteromorphic sex chromosomes can be identified from genomic data mapped to a reference genome with relative ease, because they exhibit large sex-specific differences in genomic coverage that set them apart from the autosomes. In contrast, homomorphic sex chromosomes are often extremely challenging to identify given that they exhibit few differences from each other in gene and sequence content (Palmer et al. 2019). Additionally, sex chromosomes tend to diverge in a stepwise process, and so homomorphic sex chromosomes are often a mosaic of sex-linked regions of different ages (Wright et al. 2017; Furman et al. 2020). As a result, most studies rely on a custom, combined analytical approach to identify homomorphic chromosomes, carrying out a variety of genomic analyses, each of which can provide researchers with extensive lists of candidate sex-linked regions (Palmer et al. 2019). To make this problem more challenging, each method also varies in its power to detect sex-specific sequences along the sex chromosome divergence continuum, and so, lists of candidates often do not overlap in an obvious way. For instance, sex differences in genomic coverage will identify highly divergent sex-linked regions that differ in ploidy between the X and Y, whereas sex differences in SNP density are predicted in younger regions of sex chromosomes which still retain high sequence similarity. Although it is generally understood that certain genomic approaches will be more or less effective at detecting sex chromosomes at varying stages of differentiation (Palmer et al. 2019), researchers currently lack a comprehensive workflow that incorporates approaches with the full range of sensitivities to streamline and simplify the unwieldy process of identifying sex-specific sequences in a species of interest.

#### Measures of Sex-Specific Differentiation with Coverage Approaches

In highly diverged sex chromosome systems, the X and Y (or Z and W) differ in ploidy, which will result in coverage differences between the sexes when sex-linked reads are mapped to the sex chromosomes in the reference genome (Palmer et al. 2019). The majority of the X will show approximately one half the sequencing depth in males relative to females, and vice versa for the Z chromosome, and the Y or W chromosomes will exhibit male- or female-limited coverage, respectively. This approach is often used successfully with genomic data from only a single male and single female to identify heteromorphic sex chromosomes (Palmer et al. 2019). Large-scale coverage differences have been used to identify sex chromosomes in a range of species, including insects (Vicoso and Bachtrog 2015), reptiles (Vicoso et al. 2013), birds (Sigeman, Strandh, et al. 2021), and plants (Müller et al. 2020).

#### Measures of Sex-Specific Differentiation with F_ST_

F_ST_ is an index of allelic fixation in populations, and so, if there are high levels of F_ST_ within discrete genomic regions when comparing males to females as separate populations, this would suggest that those regions differ between males and females and, therefore, recombination is absent or reduced (Zhou and Stephens 2012). However, if a sex chromosome or sex-linked region is relatively young, it is possible that sex-specific SNPs would not have time to fix completely within a population, and so would not be detected with this approach. F_ST_ has recently been used to aid in the identification of the sex determining regions in multiple stickleback and swordtail fishes (Franchini et al. 2018; Dixon et al. 2019).

#### Measures of Sex-Specific Differentiation with Genome-Wide Association Study (GWAS)

GWAS identifies associations between a phenotype and a genotype (Klein et al. 2005). By carrying out a GWAS with sex as the phenotype of interest, it is possible to identify SNPs that are strongly and weakly associated with sex. These SNPs can be fixed, or nearly fixed, in either males or females. GWAS has recently been used to identify the sex chromosomes in poplars, channel catfish (*Ictalurus punctatus*) and Atlantic salmon (*Salmo salar*) (Geraldes et al. 2015; Kijas et al. 2018; Bao et al. 2019).

#### Measures of Sex-Specific Differentiation with SNP Density

In nascent sex-linked sequences, we would expect to see differences in overall SNP density between males and females due to Y reads still mapping to the X. This is due to the divergent evolutionary trajectories of the X and Y following recombination suppression on the Y (Palmer et al 2019). SNP density can identify regions where male or female-specific SNPs are still segregating within their respective populations. A combined approach using both coverage and SNP density was able to help identify the X chromosome in guppy (*Poecilia reticulata*) (Wright et al. 2017; Sigeman, Sinclair, et al. 2021) and the switch from an XY to ZW system for white poplar (*Populus alba*) (Müller et al. 2020).

#### Measures of Sex-Specific Differentiation with Reference Genome-Free k-mer Analyses

K-mers refer to unique sequences of “k” length, that can be identified in raw sequence data. By identifying and comparing all k-mers within and between samples, all variants, including those that occur in unassembled regions can be identified and subsequently tested for associations with the phenotype of interest. This makes k-mer analyses particularly powerful, especially when working with non-model species, fragmentary genomes of low quality, or with no genome at all. This approach has been used to aid the identification of Y sequences in a range of species (Carvalho and Clark 2013; Akagi et al. 2014; Morris et al. 2018; Böhne et al. 2019). One of the drawbacks of using these methods with population-level data has been the immense computation required to calculate counts of 100s of millions or billions of k-mers across samples, however a new algorithm, kmersGWAS, that has significantly reduced compute time and storage requirements shows promise for increasing accessibility for these methods (Kokot et al. 2017; Voichek and Weigel 2020). Given its broad applicability, kmersGWAS is potentially the most powerful and innovative single method for the identification of sex-specific sequences across the entire divergence continuum using population-level sequencing data.

### The SexFindR Workflow

Given the diversity of sex chromosome systems across an evolutionary continuum that ranges from a single sex-linked SNP to highly degenerate XY, and ZW, systems, it is surprising that there is not currently a comprehensive protocol for researchers to follow in order to determine whether or not a species exhibits genetic sex determination across the full divergence gradient (fig. 1*a*). We designed SexFindR with this purpose in mind and to improve clarity and transparency across studies in the search for sex chromosomes.

**Fig. 1.**
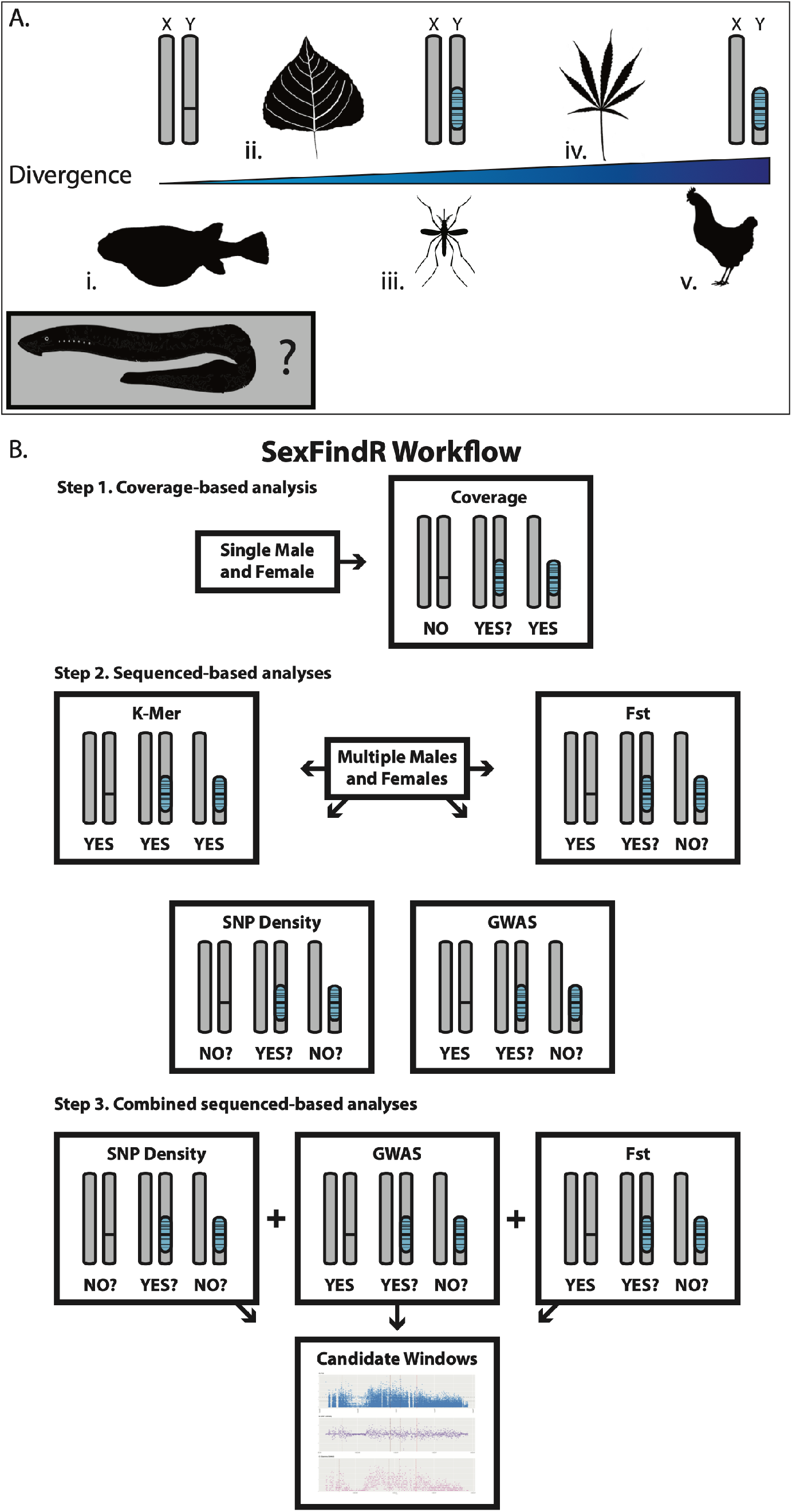
The SexFindR workflow and the sex chromosome divergence continuum. (*a*) Relative location of five selected species with varying degrees of sex chromosome divergence. i) The tiger pufferfish (*Takifugu rubripes*) has been previously shown to possess homomorphic XY chromosomes and only a single nucleotide polymorphism (SNP) perfectly linked with sex (Kamiya et al. 2012). ii) Balsam poplar (*Populus trichocarpa*) has a small, 60-100 kb, Y-like region on the left arm of chromosome 19 in the V2.2 genome (Geraldes et al. 2015; McKown et al. 2017; Müller et al. 2020). iii) The yellow fever mosquito (*Aedes aegypti*) has homomorphic sex chromosomes with a relatively small 1.3 Mb Y-like “M-locus”, but has been shown to have recombination suppression acting across approximately 40% of the 310.8 Mb chromosome on which it is contained (Fontaine et al. 2017; Aryan et al. 2020). iv) The cannabis plant (*Cannabis sativa* L) has XY chromosomes that appear nearly identical in size, but has experienced a relatively large amount of XY divergence and a small amount of Y-expansion, resulting in a Y chromosome that is ~47 Mb larger than the X chromosome (Divashuk et al. 2014; Faux et al. 2014). v) Chicken has a ZW system with a W chromosome that has degenerated in a similar manner to the mammalian Y, losing the majority of its ancestral genes (Graves 2006). Inset) The sex determination mechanism in the sea lamprey is currently unknown. (*b*) The SexFindR workflow in three steps. Each method has a schematic for three positions across the sex chromosome differentiation continuum with a YES/NO below the chromosomes to indicate whether the given method is likely to have power to detect the genomic differences associated with that sex chromosome system. Step 1. Run coverage-based analysis on a single male and female. Step 2. Run variant-based methods (k-mer, SNP density, GWAS, F_ST_) on a population genomics dataset. Step 3. Combine SNP Density, GWAS, and F_ST_ in a window analysis to filter for common hits across reference-based methods. During each step, a user has the potential to identify candidates without the need to continue to additional steps.

There are three steps in SexFindR, each of which provides an opportunity to identify sex-linked sequences in a species of interest (fig. 1*b*). Step 1: SexFindR employs a coverage-based analysis using DifCover (Smith et al. 2018), requiring only a single male and single female mapped to a reference genome, to identify large regions of chromosomal degradation or differentiation. Step 2: A variety of population genomic analyses are run in order to identify top candidate regions. These analyses include the reference-based SNP density, GWAS, and F_ST_, as well as the reference-free kmersGWAS. Although these methods have the potential to all point to the same region in the genome, they can also each identify unique regions as top candidates, which makes the robust identification of sex-linked sequences challenging. Step 3: Results from the reference-based methods are converted to 10kb windows and combined to increase the user’s power to focus on specific genomic regions. Once candidate windows are identified in Step 3, the user can return to GWAS, SNP density, and F_ST_ results to determine if fixed or nearly-fixed differences exist between the sexes in the top candidate windows.

To demonstrate the power of SexFindR, we use the workflow to identify sex-linked sequences in five species that cover the entire sex chromosome divergence continuum, from homomorphic sex chromosome systems with a single sex-linked SNP, to degenerated heteromorphic systems (fig. 1*a*). Given this broad range of applicability within a single protocol, including step-by-step methods and scripts available freely online, SexFindR represents a major improvement in the search for sex-linked sequences in non-model species. By using the method with rigid cut-offs, transparency and reproducibility can be achieved across experiments and research groups. We also apply SexFindR to a large-scale population genomics dataset we generate to assess the possibility of genetic sex determination in the sea lamprey, a species whose basis for sex determination has remained elusive for decades.

### The Search for the Mechanism of Sea Lamprey Sex Determination

Lampreys are an ancient lineage of jawless fishes comprising more than 40 extant species, approximately 500-600 million years diverged from all jawed vertebrates (Smith et al. 2013; Docker et al. 2015). This captivating phylogenetic position has led to increased study in recent years, as they can provide important information about evolution in early vertebrates (York et al. 2019). A longstanding mystery of lamprey biology is how sex is determined in this ancient vertebrate lineage, and much work over the past 50 years has tried to address this in the sea lamprey (*Petromyzon marinus*) and other lamprey species. Male-biased sex ratios under conditions of high population density or slow growth have led to suggestions of environmental sex determination (ESD) in lampreys (e.g., Torblaa and Westman 1980; Docker and Beamish 1994; Johnson et al. 2017), but evidence for ESD is equivocal (Docker et al. 2019). However, evidence for a genetic component to sex determination is lacking. The sea lamprey karyotype is complex (n=84), with many dot-like microchromosomes (Potter and Rothwell 1970; Covelo-Soto et al. 2014), and attempts to identify an X and Y, or Z and W, through karyological approaches have been unsuccessful (Ishijima et al. 2017). Additionally, Restriction-site Associated DNA sequencing (RAD-seq) was unable to identify any sex-specific sequences in European brook lamprey (*Lampetra planeri*) (Mateus et al. 2013). However, RAD-seq typically sequences only ~0.1–10% of the genome, and so, it could easily overlook sex-specific markers or even a relatively large sex-specific region.

Given the great difficulty associated with the identification of a sex-linked region or sex chromosome in sea lamprey, we believed that they would be an excellent candidate for use of the SexFindR workflow (fig. 1*a*). We carried out whole-genome resequencing of over 120 male and 120 female sea lamprey from blood and fin clips to search for the presence of fixed or nearly-fixed differences between the sexes using our new comprehensive protocol.

## New Approaches

Here, we present SexFindR, a standardized workflow for the identification of sex-linked sequences along the entire continuum of sex chromosome divergence. SexFindR incorporates the most common genomic approaches used to identify both homomorphic and heteromorphic sex-linked sequences alongside a state-of-the-art reference-free k-mer based analysis. Importantly, SexFindR combines the outputs of these analyses to screen for common candidate regions. We provide SexFindR as a 3-step protocol that allows users to identify sex-specific candidates during any of the 3 steps (fig. 1). To demonstrate the power of SexFindR, we use the workflow to identify the sex-linked sequence in five species that cover the entire divergence continuum, from homomorphic sex chromosome systems with a single sex-linked SNP, to degenerated heteromorphic systems. Finally, we apply SexFindR to our large-scale population genomics dataset to assess the possibility of genetic sex determination in the sea lamprey. The SexFindR protocol is available at the SexFindR “Read the Docs” page (https://sexfindr.readthedocs.io), and all relevant code can be found on the SexFindR GitHub (https://github.com/phil-grayson/SexFindR).

## Results

We designed the SexFindR workflow to allow researchers to robustly identify sex-specific sequences or sex chromosomes in their species of interest. We validated SexFindR in five species with known sex chromosomes that span the divergence continuum and then used it to evaluate the possibility of genetic sex determination in sea lamprey, a jawless vertebrate and invasive threat to commercial and game fisheries throughout the Laurentian Great Lakes in North America.

### SexFindR Accurately Validates Known Sex Chromosome Systems

Through the SexFindR workflow, we were able to identify the sex chromosomes in all five positive controls. In Step 1, coverage-based analyses, based on DifCover, robustly identified the known sex chromosomes in chicken, cannabis, and mosquito as outliers using only a single male and single female (fig. 1 and supplementary table S1, Supplementary Material online). This is expected as these specific exhibit heteromorphic, highly diverged sex chromosomes. Specifically, in the chicken, the Z and W chromosomes are annotated in the reference assembly. We find both exhibit significant male-biased and female-biased coverage respectively and are clear outliers relative to the rest of the genome. In cannabis, the X chromosome has not been identified but Y sequences are annotated. We were able to identify Y-linked sequences through an obvious shift in coverage relative to the rest of the genome. For mosquito, the chromosome carrying the small 1.3 Mb Y-like “M-locus” was also correctly identified as an outlier (supplementary text 1.1 and fig. S1, Supplementary Material online).

For poplar and fugu, coverage analyses in Step 1 were less effective at identifying the known sex chromosomes. While we do find that the XY sex chromosome pair in poplar is enriched for male-biased regions, and is clearly an outlier, individual regions within the chromosome show a less clear pattern. In fugu, no coverage differences are expected because only one base differs between males and females. Correspondingly, the sex chromosomes are not an outlier regarding sex differences in coverage.

We recommend stringent mapping criteria at this step, masking repetitive regions, restricting the number of mismatches when mapping reads to a reference and using only uniquely mapping reads (Palmer et al 2019). This limits incorrectly mapped reads which can mask coverage differences between the sexes and lead to the misclassification of sex-linked sequences as autosomal.

Given that the coverage-based method applied in Step 1 could not robustly identify sex chromosomes in poplar and fugu, we next applied Step 2 of SexFindR to these species. For poplar, all sequence-based methods, including the reference-free k-mer method, were able to positively identify the sex determining region as a top candidate. Importantly, all four methods independently identified the same region as outliers relative to the genome (supplementary text S1.2 and figs. S2 and S4, Supplementary Material online).

However, for fugu, results from the sequence-based analyses did not perfectly agree, although GWAS and the k-mer-based method were both able to identify the sex determining region as the most significant hits (supplementary text S1.2 and fig. S3, Supplementary Material online). Given the lack of consistent overlap between different approaches, we performed Step 3 of SexFindR on fugu. By combining the non-overlapping signals from SNP density, F_ST_, and GWAS, we were able to filter from over 35,000 10kb windows down to just 5 candidate regions, all of which are located on the previously identified sex chromosome, and one of which contained the sex-determining SNP (fig. 3 and supplementary table S2, Supplementary Material online).

### Sex-linked sequences are not detectable in the sea lamprey somatic genome

Given our successful identification of all known sex chromosomes in species that span the sex chromosome divergence continuum, from a single SNP to highly degenerate systems, we next applied SexFindR to sea lamprey, a species with an unknown and elusive sex determination system.

First, coverage-based methods in Step 1 did not find consistent male-female enrichment, indicating that lamprey does not have a large region of recombination suppression or a degenerated, or expanded, Y or W chromosome. However, while the lamprey does not have any obvious DifCover regions that exhibit unusual sex differences in coverage (fig. 2*f* left plot and supplementary table S1, Supplementary Materials online), there is a single scaffold that is enriched for sex-biased regions relative to the rest of the genome (fig. 2*f* right plot). This possible sex-linked candidate was disproven with additional analyses, and outliers in the depth analysis appear to be due to large segregating deletions within the sea lamprey (supplementary fig. S7 and text S2.1, Supplementary Materials online).

**Fig. 2.**
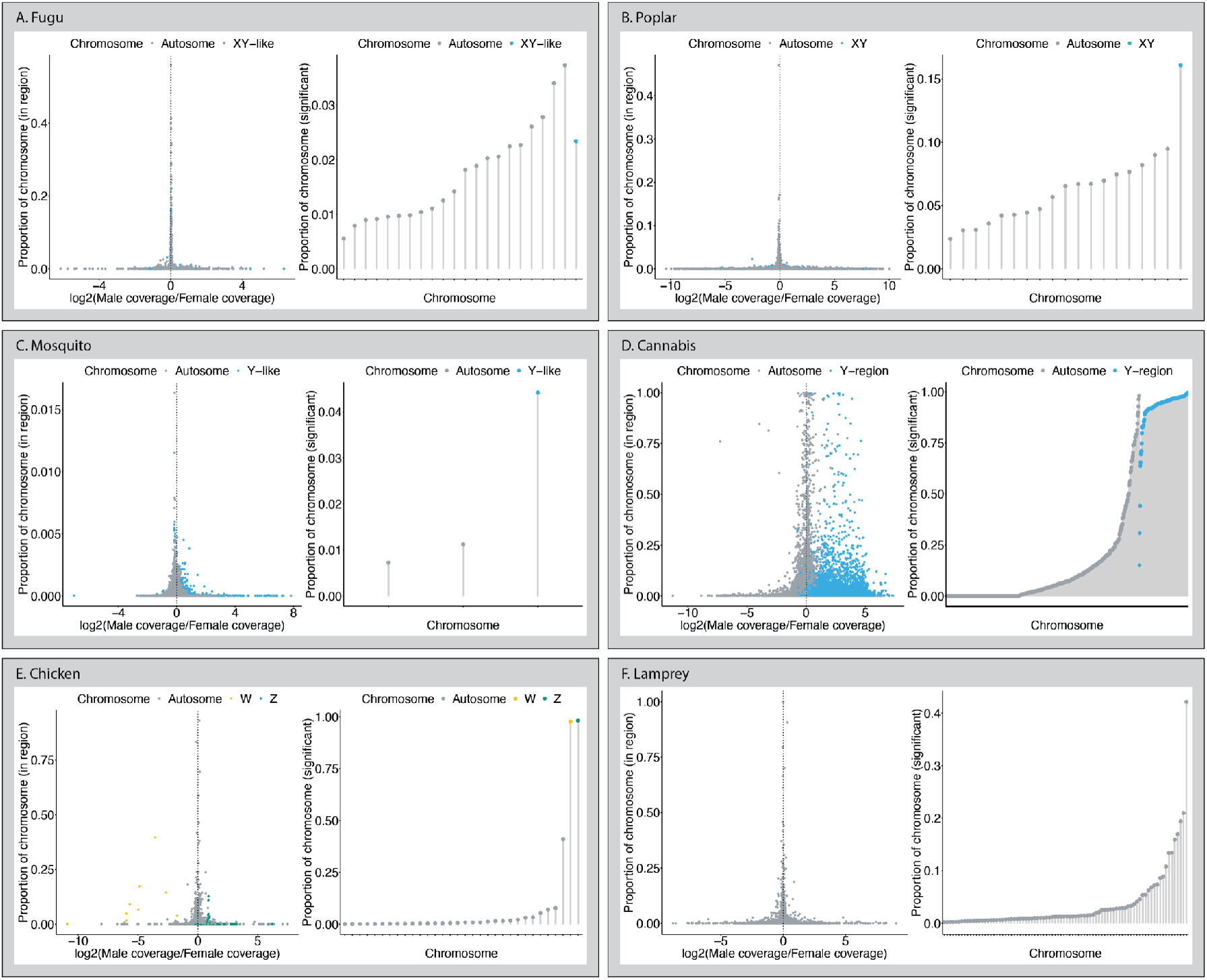
Sex differences in coverage across the genome calculated using DifCover, which breaks chromosomes (or scaffolds) into regions with similar coverage differences between the sexes. For each species (A-F), the left plot has log2(male / female coverage difference) on the X axis and the proportion of the full chromosome or scaffold that specific region occupies on the Y axis. The right plot collects all significant regions for each chromosome or scaffold and plots the total proportion of the chromosomal length that displays significantly different coverage between males and females. Known sex chromosomes are shown in blue (X or Y), orange (W), or green (Z) for each species, except for the lamprey for which has an unknown sex determination system.

**Fig. 3.**
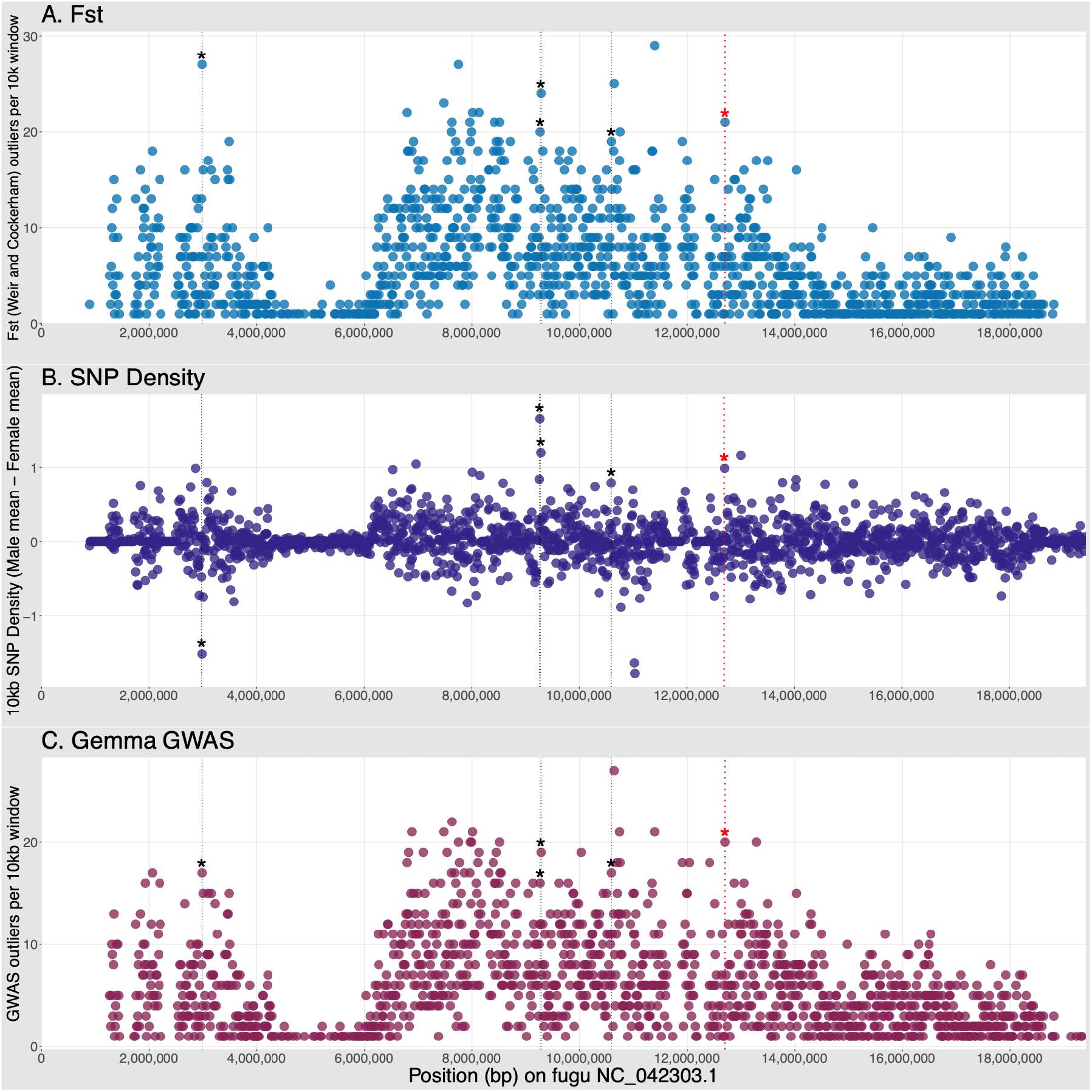
SexFindR Step 3 combined results for (*a*) F_ST_, (*b*) SNP density, and (*c*) GWAS identified the fugu sex determining region (red dotted line and red asterisk), alongside four additional candidate windows (thin black dotted lines with black asterisks), all on the known sex chromosome (NC_042303.1).

Next, we found no fixed (or nearly-fixed) sequence differences between male and female sea lamprey in Step 2 using SNP density, GWAS, or F_ST_, as well as the reference-free kmersGWAS (supplementary text S1.2, Supplementary Materials online). Using the combined approach in Step 3, we did identify a small number of 10kb candidate windows (supplementary table S3, Supplementary Materials online), but results were consistently less significant than those from fugu, and when these regions were examined in detail, it was determined that the SNPs driving the signal were segregating in both males and females (supplementary text S1.3, Supplementary Materials online). Notably, we ran SexFindR on lamprey samples collected across a number of different populations. It is possible that the sex determination mechanism or degree of sex chromosome divergence varies across populations, as has been observed in a number of fish, amphibian and reptile species (Furman et al 2020). We tested for this by analysing data from each of the 9 populations independently but found no consistent sex-specific SNPs (supplementary text S2.2, Supplementary Materials online). Given that fugu only has a single fixed difference between males and females, and that we were able to identify that site with the SexFindR workflow, this lends support to our conclusion that sea lamprey do not carry sex-linked sequences in their somatic genome.

## Discussion

In this study, we designed a comprehensive workflow, SexFindR, for using next-generation short-read sequencing data to investigate the signals of genetic sex determination in a species of interest. We validate this method in five species with known sex determination systems that span the continuum from a single sex-linked SNP to heteromorphic sex chromosomes. Different approaches to detect sex chromosomes possess varying levels of power along the sex chromosome continuum, and the data and results are often very noisy and difficult to interpret. SexFindR provides a robust and standardised workflow for the community to identify sex-linked sequences. In doing so, we anticipate that this will reduce the potential for discrepancies between studies arising from differences in methodology or parameter choice and facilitate the identification of sex chromosomes in a wide range of non-model species.

Importantly, we used SexFindR to investigate the possibility of genetic sex determination in the sea lamprey, a jawless fish whose sex determination system has remained elusive for decades. Proving a negative in genomics is quite challenging and publishing a negative result even more so. Without a unified and transparent pipeline, such as SexFindR, it is unclear if sex-linked sequences have not been found for a given species because the right method with the correct parameters has yet to be applied, or if the species truly does not possess sex-linked sequences within the data analyzed. The lack of a consistent or compelling signal of a sex-linked sequence within the sea lamprey genome strongly suggests that there are no fixed (or nearly fixed) sex-specific sequences within our dataset. Although this result does not unequivocally solve the question of sex determination in sea lamprey, it does provide further support for mechanisms other than the standard XY or ZW systems with sex-linked sequence.

It is possible that sea lamprey exhibit entirely environmental sex determination. Environmental sex determination has been suggested for some lamprey species, but these data are inconclusive due to the difficulty associated with studying the long and complex lamprey life cycle (Docker et al. 2019). If, on the other hand, sea lamprey do possess a fixed genetic basis for phenotypic sex, it is possible that this region is not present in the somatic sequences. It has been shown that the sea lamprey undergo a programmed genome rearrangement (PGR) early in the first days of embryonic development (Smith et al. 2009; Smith et al. 2010; Smith et al. 2013; Smith et al. 2018). This PGR results in the deletion of approximately 20% of the germline genome (~500,000 bp) from all somatic cells. In sea lamprey, the ovary and testis develop at different times, both years after this PGR, but it is possible that sex-specific germline sequence could be responsible for this differentiation (Docker et al. 2019). Despite our use of a germline reference genome, the samples used in this study all originated from a large population genomics project which collected blood and fin clips, which are somatic tissues; therefore, any sex-specific sequence present only in the germline genome would not be identified within this dataset. Carrying out population genomics using the lamprey germ cells, or examination of the transcriptome or regulatory landscape in developing gonads could shed light on this possibility. In fact, recent work carried out in our laboratory using transcriptomics on developing gonadal tissue from sea lamprey has uncovered a striking signal (Yasmin et al. 2021). By conducting comparative transcriptomic experiments on developing testes and ovaries, Yasmin and colleagues (2021) were able to discover that the 638 genes found within the germline-specific regions exhibit a 36x greater odds of being expressed in testes compared to ovaries, while many of these genes have paralogs in the somatic genome that do not experience differential expression. These male-specific germline genes are likely to be involved in gonadal differentiation and possibly sex determination itself, and include genes known to be important for sex determination in other species.

## Materials and Methods

### Positive Controls

#### SexFindR Step 1. Coverage-Based Analysis

Samples for fugu, poplar, mosquito, cannabis and chicken were downloaded from various sources and mapped to their reference genomes (supplementary table S4, Supplementary Materials online) using Bowtie2 (v 2.3.4.3) (Langmead and Salzberg 2013). Using stringent mapping threshold, such as uniquely mapping reads, is critical to minimise noise and ensure that coverage differences between males and females can be detected (Darolti et al. 2021) BAM files were sorted with Samtools sort and indexed using Samtools index (both v 1.10).

Given that in an XY or ZW system the homogametic sex has two copies of the X or Z, the heterogametic sex, XY males and ZW females, should show approximately one half the sequencing depth for any reads mapping to the X or Z respectively. With sequence data from a single male and single female, large regions of sequence divergence or chromosomal degeneration can be identified if present. To test for genomic regions that differ in coverage between males and females, we used DifCover, a program that searches for differences in coverage via an interval-based method that has been used to identify both germline-specific and sex-specific markers (Keinath et al. 2018; Smith et al. 2018; Timoshevskiy et al. 2019). Unlike some window-based approaches, DifCover does not report scores for each window of a predefined length. Instead, it carries out a two-step process: first, given a user-defined length of valid bases (e.g., 1000 bp), DifCover scans each chromosome and scaffold to generate initial windows, where valid bases fall within a range of sequencing depths (e.g., 10x to 240x) in at least one of the samples analyzed. Once average male/female coverage differences have been calculated for these initial windows, these windows are merged based on their similarity to surrounding windows to generate the variably-sized regions we report. The goal with the “valid bases” is to eliminate the signal from problematic regions, including highly repetitive and gapped alignments (Smith et al. 2018).

Previously, DifCover has been run with a log2 cut off of 2, which represents a 4-fold change in mapping between the two samples. Given that the expectation for ZW and XY systems should be a change lower than 4-fold, DifCover was run with a lower log2 cut off of 0.740, which corresponds to a ratio of 1.2/2 or 2/1.2 in case the standard cut off of 4 was too high to identify a simple 2-fold change in coverage expected for the X in XY systems and the Z in ZW systems. This cut off allows us to identify Y- or W-linked reads that show very high levels of enrichment in the male or female respectively and also identify X- or Z-linked reads that show a ratio closer to 1/2 or 2/1 between males and females. All other parameters remained at default settings.

#### SexFindR Step 2. Variant-Based Analysis

For the positive controls, Step 2 of SexFindR was only run on poplar and fugu, given that Step 1 was able to correctly identify sex chromosomes in mosquito, cannabis and chicken. Population genomic samples for fugu and poplar were downloaded from NCBI (supplementary tables S4-S6, Supplementary Materials online). For reference-based analyses (e.g., F_ST_, GWAS, and SNP density), reads were mapped to the reference genome using Bowtie2 as above. BAM files were used as input to the Platypus Variant Caller (https://github.com/andyrimmer/Platypus) using joint calling (Rimmer et al. 2014). The resulting VCF was filtered to retain only those sites that had “PASS” under the “FILTER” column using bcftools (v 1.9) (Li et al. 2009).

For F_ST_, the VCF was also filtered to include only biallelic sites with no missing data. F_ST_ was calculated, comparing the males and females as separate populations at each SNP using VCFtools (v0.1.14) (Danecek et al. 2011). F_ST_ sites were ranked from highest to lowest F_ST_ value to identify top candidates.

For GWAS, the VCF was used as input for the program GEMMA (v0.98.1), to identify any SNPs that correlate significantly with phenotypic sex (Zhou and Stephens 2012). VCFtools was used to convert the VCF file to PLINK format while removing indels, requiring at least 50% coverage of the site, and requiring the minor allele frequency to be between 0.05 and 0.95. GEMMA was run with default parameters for a binary trait (affected/unaffected), which automatically filters for biallelic sites. Sites were sorted from lowest to highest p_lrt to identify top candidates.

For SNP density, the VCF file was filtered to include only binary sites and then a single VCF file was created for each individual sample using bcftools. VCFtools snpdensity was used to calculate SNP density for each sample in non-overlapping 10kb windows across the genome. The mean SNP density for males and females and the difference between the population means were calculated using an in-house R script (see Read the Docs). In order to determine if these differences were greater than expected by chance, p-values were assigned to the difference observed in each window using a permutation test. For each permutation, the sex of all samples was randomized while maintaining the correct number of males and females in the dataset. An average SNP density for these permutated “males” and “females” was calculated as above. Following the completion of 100,000 permutations, a p-value was calculated for each window in the genome, based on how many permutations were more extreme than the true calculated value (either a greater positive or negative difference between the permuted averages compared to the true average). Windows with p <= 0.05 were retained, sorted by p-value, and finally sorted by absolute difference in change between the male and female SNP density means to identify top candidates.

For kmersGWAS, we used default methodology with slight modifications. For each sample, 31-mers were counted from unmapped fastq data as canonical (count has a minimum of 2), and noncanonical (no minimum count). The two k-mer lists were combined with strand information to generate a single file per sample. The k-mers for all samples were combined using the following criteria: the k-mer has to appear in at least 5 individuals and it must appear in the canonized list for at least 20% of the individuals that it appears in. A presence/absence count table was generated for samples for every k-mer that passed the filter from the previous step. The k-mer table was converted to PLINK binary format with a minor allele frequency of 0.05 and minor allele count of five. PLINK 1.07 was run with the --noweb, --bfile, and --allow-no-sex flags to generate p-values for every k-mer in the analysis. k-mers were sorted by p-value and the top k-mers were filtered to a top candidate out-file. The top candidate out-file was parsed using plink_to_abyss_kmers.py to pass to ABYSS (Jackman et al. 2017), which assembles the k-mers into contigs (Jackman et al. 2017). Resulting contigs were used as queries for blastn on NCBI to validate the candidates and determine if unassembled sequences were highly significant.

#### SexFindR Step 3. Combined Variant-Based Analysis

For the positive controls, Step 3 was only run on fugu, since Step 2 successfully identified the known poplar sex determining region. We developed an in-house linux/R/python workflow to filter for top candidates based on data from F_ST_, SNP density, and GWAS analyses. This produces a single R data frame where 10kb windows are ranked for each analysis to leverage the power of these varied approaches and identify windows that are identified as outliers in multiple analyses. The theoretical inspiration for carrying out this analysis come from the Composite of Multiple Signals (CMS) program (Grossman et al. 2010).

With SNP density already partitioned into 10kb windows and ranked, we selected the top 5% of SNPs from the previously-ranked GWAS and F_ST_ outputs, and counted the number of sites within those same 10kb windows for both GWAS and F_ST_. Windows were then ranked for each analysis based on the total number of sites contained within. Candidate windows were considered if they ranked within the top 100 for SNP density, GWAS, and F_ST_.

### Sea Lamprey

#### Sample Collection

Two sets of lamprey samples are used in this study. First, adult sea lamprey (n = 20, supplementary table S7, Supplementary Materials online) were collected in June 2019 in the Cheboygan River, MI, USA, a tributary of Lake Huron, during their upstream (spawning) migration. Sterile gloves, pipettes, scalpels, parafilm, and collection dishes were used for each lamprey, and males and females were collected on separate days to control for cross-sample contamination. Blood was collected into an EDTA-coated Vacutainer tube. For the second set of samples, additional whole upstream-migrating sea lamprey (n = 246, supplementary table S8, Supplementary Materials online) were collected between May and August 2019 for a large-scale population genomics project that covered the Finger Lakes in New York state, all five Great Lakes (Erie, Huron, Michigan, Ontario, and Superior), and an anadromous population from the Connecticut River (at the Holyoke Dam), MA, USA. Lamprey from the initial Huron samples will be referred to as the “original Huron samples” whenever there is a need to distinguish them from the large population dataset.

Abbreviated Protocols for Minimal Animal Involvement, one for the adult sea lamprey collected in June 2019 in the Cheboygan River and one for those collected May–August 2019 from the Great Lakes, Finger Lakes, and Connecticut River, were approved by the Fort Garry Campus Animal Care Committee at the University of Manitoba. Sea lamprey were captured at traps during their spawning migration as part of the ongoing sea lamprey control efforts in the Great Lakes basin or collected for other projects, and they were euthanized by Sea Lamprey Control personnel or colleagues from the U.S. Geological Survey according to their approved protocols.

#### Sea Lamprey Short-Read DNA Extraction, Sequencing, and Preprocessing

For the original Huron samples, DNA was extracted from blood using the DNeasy Blood & Tissue kit with minor modifications to the standard protocol. 200 uL of blood was used for each individual and the final elution was carried out in 50 uL of Buffer AE. Following quality control and quantification, DNA was shipped on ice to The Center for Applied Genomics (TCAG) where libraries were prepared and sequenced. DNA was first quantified with the Qubit dsDNA HS assay and sheared to a standard 400-bp on the Covaris LE220. Libraries were generated using the Illumina TruSeq PCR-free library preparation protocol with 400-bp insert size on an Agilent Bravo automation system (model B). 700 ng of starting material was used for each library, except for M_9d and M12_d, which only had 535 ng and 615 ng of DNA respectively. qPCR was performed on the resulting libraries and they were multiplexed into two pools. Pool1 contained 16 samples (low coverage of ~15x for 8 males and 8 females) and Pool2 contained 4 samples (high coverage of ~60x for 2 males and 2 females). Each pool was sequenced across 3 flowcells of a single Illumina HiSeq X run with 150-bp paired-end reads. Initially, lamprey had been visually sexed at the time of collection. This resulted in one female being mis-labeled as a male (M2_d in all downstream analyses). Following this discovery, all lamprey carcasses were thawed for dissection to visually identify either a testis or ovary. An additional flowcell of Illumina HiSeq X 150-bp paired-end sequencing was carried out for M8_d to replace M2_d as a high coverage male. For the large population samples, DNA was extracted from fin clips taken from sexed frozen carcasses using standard protocol from Qiagen’s DNeasy Blood & Tissue kit with minor modifications. The tissue was allowed to incubate for 18-20 hours for full proteinase K digestion, and 100 uL of warmed (70°C) Buffer AE was allowed to incubate for 1 minute prior to final elution. Following DNA extraction, all samples were once again sent to TCAG for identical library preparation, but the resulting libraries were sequenced on an Illumina NovaSeq S4 flowcell at 2×150 bp. FastQC (v 0.11.8) was run on all libraries as a quality control measure, and all fastq files performed as expected (Andrews 2010). Over-represented sequences and adapters were removed from raw sequencing data using Trimmomatic (v 0.36) (Bolger et al. 2014) prior to second run of FastQC to confirm trimming.

#### Sea Lamprey Read Mapping and Variant Calling

Following quality control, sea lamprey reads were mapped to the newest germline lamprey genome (kPetMar1 from NCBI - https://ftp.ncbi.nlm.nih.gov/genomes/all/GCF/010/993/605/GCF_010993605.1_kPetMar1.pri/GCF_010993605.1_kPetMar1.pri_genomic.fna.gz) using Bowtie2 (v 2.3.4.3) with stringent mapping criteria (-X 1000 --fr --no-mixed --no-discordant) (Langmead and Salzberg 2013). BAM files were sorted with Samtools sort, and indexed using Samtools index (both v 1.10) (Li et al. 2009). Deduplication was carried out using Picard MarkDuplicates (v 2.20.6) and the final BAM files were sorted and indexed as above. The BAM files were used as input to the Platypus Variant Caller (https://github.com/andyrimmer/Platypus) using all samples at once in order to call SNPs jointly (Rimmer et al. 2014). The resulting VCF was filtered to retain only those sites that had “PASS” under the “FILTER” column using bcftools (v 1.9) (Li et al. 2009).

#### SexFindR Step 1. Coverage-Based Analysis

Following deduplication, a single male and single female lamprey with deep sequencing ~60x were selected for DifCover analysis, which was conducted as above. To further screen the resulting candidate regions, a 2-by-2 DifCover analysis was also carried out (supplemental text 2.1, Supplementary Materials online).

For lamprey, additional runs of DifCover were conducted to account for our sequencing strategy which included deep sequencing for 2 males and 2 females. Each deeply sequenced female was run against each deeply sequenced male, resulting in 4 experimental comparisons. The two males were run against each other, as were the two females, to serve as controls. Finally, all low-coverage females from the original Huron samples dataset were merged to produce a single file which was compared to a merged file containing all low-coverage males. Bedtools (v2.25.0) makewindows was used to generate 10kb windows across the genome, and bedtools annotate (v2.25.0) was used to map significant regions back to the genome. In R, filtering was carried out to identify candidate regions of overlap in all 4 experimental comparisons that were not found in the control runs. These regions were then compared against the regions identified in the low-coverage DifCover run to search for consistent regions with differential coverage between the males and females.

#### SexFindR Step 2. Sequenced-Based Analysis

All steps were carried out as described in the Positive Controls section for F_ST_, GWAS, SNP density and kmersGWAS.

#### SexFindR Step 3. Combined Sequence-Based Analyses

All steps were carried out as described in the Positive Controls section to combine the results from F_ST_, GWAS and SNP density.

## Acknowledgments

Dr. Nick Johnson (US Geological Survey, USGS), Matt Symbal (US Fish and Wildlife, Great Lakes), Gale Bravener (Fisheries and Oceans Canada, Great Lakes), Dr. Ted Castro-Santos (USGS, Holyoke Dam, Connecticut River), Emily Zollweg-Horan (NY State Department of Environmental Conservation, Cayuga Lake), and Brad Hammers (NY State Department of Environmental Conservation, Seneca Lake) for the sea lamprey samples, and Arfa Khan (University of Manitoba) for performing the DNA extractions.

This research was funded by the Great Lakes Fishery Commission Sea Lamprey Research Program (2018_DOC_54073).

## Author Contribution

MD, CG, AW, and PG conceived the study. PG developed SexFindR, performed the analysis, and wrote the paper with input from all the authors.

## Data Availability

All relevant code can be found at the SexFindR GitHub repo (https://github.com/phil-grayson/SexFindR) and a detailed walkthrough of the SexFindR protocol can be found at the SexFindR “Read the Docs” page (https://sexfindr.readthedocs.io/en/latest/). Raw sequencing data for sea lamprey will be deposited at NCBI’s SRA prior to publication.

## Notes

https://github.com/phil-grayson/SexFindR

https://sexfindr.readthedocs.io

### Competing Interest Statement

The authors have declared no competing interest.

## References

Akagi T, Henry IM, Tao R, Comai L. 2014. A y-chromosome-encoded small RNA acts as a sex determinant in persimmons. Science (80-). 346(6209):646–650. doi:10.1126/science.1257225.

Andrews S. 2010. FastQC: A Quality Control Tool for High Throughput Sequence Data. https://github.com/s-andrews/FastQC.

Aryan A, Anderson MAE, Biedler JK, Qi Y, Overcash JM, Naumenko AN, Sharakhova M V., Mao C, Adelman ZN, Tu Z. 2020. Nix alone is sufficient to convert female Aedes aegypti into fertile males and myo-sex is needed for male flight. Proc Natl Acad Sci U S A. 117(30):17702–17709. doi:10.1073/pnas.2001132117.

Bao L, Tian C, Liu S, Zhang Y, Elaswad A, Yuan Z, Khalil K, Sun F, Yang Y, Zhou T, et al. 2019. The Y chromosome sequence of the channel catfish suggests novel sex determination mechanisms in teleost fish. BMC Biol. 17(1):1–16. doi:10.1186/s12915-019-0627-7.

Böhne A, Weber AAT, Rajkov J, Rechsteiner M, Riss A, Egger B, Salzburger W. 2019. Repeated evolution versus common ancestry: Sex chromosome evolution in the haplochromine CICHLIDX Pseudocrenilabrus philander. Genome Biol Evol. 11(2):439–458. doi:10.1093/gbe/evz003.

Bolger AM, Lohse M, Usadel B. 2014. Trimmomatic: A flexible trimmer for Illumina sequence data. Bioinformatics. 30(15):2114–2120. doi:10.1093/bioinformatics/btu170.

Carvalho AB, Clark AG. 2013. Efficient identification of Y chromosome sequences in the human and Drosophila genomes. Genome Res. 23(11):1894–1907. doi:10.1101/gr.156034.113.

Charlesworth D, Charlesworth B, Marais G. 2005. Steps in the evolution of heteromorphic sex chromosomes. Heredity (Edinb). 95(2):118–128. doi:10.1038/sj.hdy.6800697.

Covelo-Soto L, Morán P, Pasantes JJ, Pérez-García C. 2014. Cytogenetic evidences of genome rearrangement and differential epigenetic chromatin modification in the sea lamprey (Petromyzon marinus). Genetica. 142(6):545–554. doi:10.1007/s10709-014-9802-5.

Danecek P, Auton A, Abecasis G, Albers CA, Banks E, DePristo MA, Handsaker RE, Lunter G, Marth GT, Sherry ST, et al. 2011. The variant call format and VCFtools. Bioinformatics. 27(15):2156–2158. doi:10.1093/bioinformatics/btr330.

Darolti I, Almeida P, Wright AE, Mank JE. 2021. A comparison of methodological approaches to the study of young sex chromosomes: A case study in Poecilia. bioRxiv.:2021.11.29.470452. doi:10.1101/2021.11.29.470452. http://biorxiv.org/content/early/2021/12/01/2021.11.29.470452.abstract.

Divashuk MG, Alexandrov O S, Razumova OV., Kirov I V., Karlov GI. 2014. Molecular cytogenetic characterization of the dioecious Cannabis sativa with an XY chromosome sex determination system. PLoS One. 9(1):1–7. doi:10.1371/journal.pone.0085118.

Dixon G, Kitano J, Kirkpatrick M. 2019. The Origin of a New Sex Chromosome by Introgression between Two Stickleback Fishes. Mol Biol Evol. 36(1):28–38. doi:10.1093/molbev/msy181.

Docker MF, Beamish FWH. 1994. Age, growth, and sex ratio among populations of least brook lamprey, Lampetra aepyptera, larvae: an argument for environmental sex determination. In: Balon EK, Bruton MN, Noakes DLG, editors. Women in ichthyology: an anthology in honour of ET, Ro and Genie. Dordrecht: Springer Netherlands. p. 191–205. https://doi.org/10.1007/978-94-011-0199-8_16.

Docker MF, Beamish FWH, Yasmin T, Bryan MB, Khan A. 2019. The Lamprey Gonad. In: Lampreys: Biology, Conservation and Control, Vol. 2. Springer, Dordrecht. p. 1–186. http://dx.doi.org/10.1007/978-94-024-1684-8_1.

Docker MF, Hume JB, Clemens BJ. 2015. Introduction: a surfeit of lampreys. In: Lampreys: biology, conservation and control. Springer, Dordrecht. p. 1–34.

Faux AM, Berhin A, Dauguet N, Bertin P. 2014. Sex chromosomes and quantitative sex expression in monoecious hemp (Cannabis sativa L.). Euphytica. 196(2):183–197. doi:10.1007/s10681-013-1023-y.

Feron R, Pan Q, Wen M, Imarazene B, Jouanno E, Anderson J, Herpin A, Journot L, Parrinello H, Klopp C, et al. 2021. RADSex: A computational workflow to study sex determination using restriction site-associated DNA sequencing data. Mol Ecol Resour. 21(5):1715–1731. doi:10.1111/1755-0998.13360.

Fontaine A, Filipović I, Fansiri T, Hoffmann AA, Cheng C, Kirkpatrick M, Rašić G, Lambrechts L. 2017. Extensive genetic differentiation between homomorphic sex chromosomes in the mosquito vector, Aedes aegypti. Genome Biol Evol. 9(9):2322–2335. doi:10.1093/gbe/evx171.

Franchini P, Jones JC, Xiong P, Kneitz S, Gompert Z, Warren WC, Walter RB, Meyer A, Schartl M. 2018. Long-term experimental hybridisation results in the evolution of a new sex chromosome in swordtail fish. Nat Commun. 9(1):1–11. doi:10.1038/s41467-018-07648-2. http://dx.doi.org/10.1038/s41467-018-07648-2.

Furman BLS, Metzger DCH, Darolti I, Wright AE, Sandkam BA, Almeida P, Shu JJ, Mank JE, Fraser B. 2020. Sex Chromosome Evolution: So Many Exceptions to the Rules. Genome Biol Evol. 12(6):750–763. doi:10.1093/gbe/evaa081.

Gautier M. 2014. Using genotyping data to assign markers to their chromosome type and to infer the sex of individuals: a Bayesian model-based classifier. Mol Ecol Resour. 14(6): 1141–1159. doi:10.1111/1755-0998.12264.

Geraldes A, Hefer CA, Capron A, Kolosova N, Martinez-Nuñez F, Soolanayakanahally RY, Stanton B, Guy RD, Mansfield SD, Douglas CJ, et al. 2015. Recent y chromosome divergence despite ancient origin of dioecy in poplars (Populus). Mol Ecol. 24(13):3243–3256. doi:10.1111/mec.13126.

Graves JAM. 2006. Sex chromosome specialization and degeneration in mammals. Cell. 124(5):901–914. doi:10.1016/j.cell.2006.02.024.

Grossman SR, Shylakhter I, Karlsson EK, Byrne EH, Morales S, Frieden G, Hostetter E, Angelino E, Garber M, Zuk O, et al. 2010. A composite of multiple signals distinguishes cuasal variants in regions of positive selection. Science (80-). 166(February):2008–2011.

Ishijima J, Uno Y, Nunome M, Nishida C, Kuraku S, Matsuda Y. 2017. Molecular cytogenetic characterization of chromosome site-specific repetitive sequences in the Arctic lamprey (Lethenteron camtschaticum, Petromyzontidae). DNA Res. 24(1):93–101. doi:10.1093/dnares/dsw053.

Jackman SD, Vandervalk BP, Mohamadi H, Chu J, Yeo S, Hammond SA, Jahesh G, Khan H, Coombe L, Warren RL, et al. 2017. ABySS 2.0: Resource-Efficient Assembly of Large Genomes using a Bloom Filter Effect of Bloom Filter False Positive Rate. Genome Res. 27:768–777. doi:10.1101/gr.214346.116.Freely.

Johnson NS, Swink WD, Brenden TO. 2017. Field study suggests that sex determination in sea lamprey is directly influenced by larval growth rate. Proc R Soc B Biol Sci. 284(1851). doi:10.1098/rspb.2017.0262.

Kamiya T, Kai W, Tasumi S, Oka A, Matsunaga T, Mizuno N, Fujita M, Suetake H, Suzuki S, Hosoya S, et al. 2012. A trans-species missense SNP in Amhr2 is associated with sex determination in the tiger Pufferfish, Takifugu rubripes (Fugu). PLoS Genet. 8(7). doi:10.1371/journal.pgen.1002798.

Keinath MC, Timoshevskaya N, Timoshevskiy VA, Voss SR, Smith JJ. 2018. Miniscule differences between sex chromosomes in the giant genome of a salamander. Sci Rep. 8(1): 1–14. doi:10.1038/s41598-018-36209-2.

Kijas J, McWilliam S, Naval Sanchez M, Kube P, King H, Evans B, Nome T, Lien S, Verbyla K. 2018. Evolution of Sex Determination Loci in Atlantic Salmon. Sci Rep. 8(1):1–11. doi:10.1038/s41598-018-23984-1.

Klein RJ, Zeiss C, Chew EY, Tsai JY, Sackler RS, Haynes C, Henning AK, SanGiovanni JP, Mane SM, Mayne ST, et al. 2005. Complement factor H polymorphism in age-related macular degeneration. Science. 308(5720):385–389. doi:10.1126/science.1109557.

Kokot M, Dlugosz M, Deorowicz S. 2017. KMC 3: counting and manipulating k-mer statistics. Bioinformatics. 33(17):2759–2761. doi:10.1093/bioinformatics/btx304.

Langmead B, Salzberg S. 2013. Bowtie2. Nat Methods. 9(4):357–359. doi:10.1038/nmeth.1923.Fast. https://www.ncbi.nlm.nih.gov/pmc/articles/PMC3322381/pdf/nihms-366740.pdf.

Li H, Handsaker B, Wysoker A, Fennell T, Ruan J, Homer N, Marth G, Abecasis G, Durbin R, 1000 Genome Project Data Processing Subgroup. 2009. The Sequence Alignment Map format and SAMtools. Bioinformatics. 25(16):2078–2079. doi:10.1093/bioinformatics/btp352.

Mateus CS, Stange M, Berner D, Roesti M, Quintella BR, Alves MJ, Almeida PR, Salzburger W. 2013. Strong genome-wide divergence between sympatric European river and brook lampreys. Curr Biol. 23(15):R649–R650. doi:10.1016/j.cub.2013.06.026. http://dx.doi.org/10.1016/j.cub.2013.06.026.

McKown AD, Klápště J, Guy RD, Soolanayakanahally RY, La Mantia J, Porth I, Skyba O, Unda F, Douglas CJ, El-Kassaby YA, et al. 2017. Sexual homomorphism in dioecious trees: Extensive tests fail to detect sexual dimorphism in Populus. Sci Rep. 7(1):1–14. doi:10.1038/s41598-017-01893-z.

Morris J, Darolti I, Bloch NI, Wright AE, Mank JE. 2018. Shared and species-specific patterns of nascent Y chromosome evolution in two guppy species. Genes (Basel). 9(5): 11–14. doi:10.3390/genes9050238.

Müller NA, Kersten B, Leite Montalvão AP, Mähler N, Bernhardsson C, Bräutigam K, Carracedo Lorenzo Z, Hoenicka H, Kumar V, Mader M, et al. 2020. A single gene underlies the dynamic evolution of poplar sex determination. Nat Plants. 6(6):630–637. doi:10.1038/s41477-020-0672-9. http://dx.doi.org/10.1038/s41477-020-0672-9.

Muyle A, Käfer J, Zemp N, Mousset S, Picard F, Marais GAB. 2016. Sex-detector: A probabilistic approach to study sex chromosomes in non-model organisms. Genome Biol Evol. 8(8):2530–2543. doi:10.1093/gbe/evw172.

Palmer DH, Rogers TF, Dean R, Wright AE. 2019. How to identify sex chromosomes and their turnover. Mol Ecol. 28(21):4709–4724. doi:10.1111/mec.15245.

Potter IC, Rothwell B. 1970. The mitotic chromosomes of the lamprey, Petromyzon marinus L. Experientia. 26(4):429–430. doi:10.1007/BF01896930.

Rangavittal S, Stopa N, Tomaszkiewicz M, Sahlin K, Makova KD, Medvedev P. 2019. DiscoverY: A classifier for identifying y chromosome sequences in male assemblies. BMC Genomics. 20(1):1–11. doi:10.1186/s12864-019-5996-3.

Rimmer A, Phan H, Mathieson I, Iqbal Z, Twigg SRF, Wilkie AOM, Mcvean G, Lunter G. 2014. Integrating mapping-, assembly-and haplotype-based approaches for calling variants in clinical sequencing applications. Nat Genet. 46(8):912–918. doi:10.1038/ng.3036.

Sigeman H, Sinclair B, Hansson B. 2021. FindZX: an automated pipeline for detecting and visualising sex chromosomes using whole-genome sequencing data.:1–22.

Sigeman H, Strandh M, Proux-Wéra E, Kutschera VE, Ponnikas S, Zhang H, Lundberg M, Soler L, Bunikis I, Tarka M, et al. 2021. Avian Neo-Sex Chromosomes Reveal Dynamics of Recombination Suppression and W Degeneration. Mol Biol Evol. 38(12):5275–5291. doi:10.1093/molbev/msab277.

Smith JJ, Antonacci F, Eichler EE, Amemiy CT. 2009. Programmed loss of millions of base pairs from a vertebrate genome. Proc Natl Acad Sci U S A. 106(27): 11212–11217. doi:10.1073/pnas.0902358106.

Smith JJ, Kuraku S, Holt C, Sauka-Spengler T, Jiang N, Campbell MS, Yandell MD, Manousaki T, Meyer A, Bloom OE, et al. 2013. Sequencing of the sea lamprey (Petromyzon marinus) genome provides insights into vertebrate evolution. Nat Genet. 45(4):415–421. doi:10.1038/ng.2568. http://dx.doi.org/10.1038/ng.2568.

Smith JJ, Stuart AB, Sauka-Spengler T, Clifton SW, Amemiya CT. 2010. Development and analysis of a germline BAC resource for the sea lamprey, a vertebrate that undergoes substantial chromatin diminution. Chromosoma. 119(4):381–389. doi:10.1007/s00412-010-0263-z.

Smith JJ, Timoshevskaya N, Ye C, Holt C, Keinath MC, Parker HJ, Cook ME, Hess JE, Narum SR, Lamanna F, et al. 2018. The sea lamprey germline genome provides insights into programmed genome rearrangement and vertebrate evolution. Nat Genet. 50(2):270–277. doi:10.1038/s41588-017-0036-1. http://dx.doi.org/10.1038/s41588-017-0036-1.

Timoshevskiy VA, Timoshevskaya NY, Smith JJ. 2019. Germline-Specific Repetitive Elements in Programmatically Eliminated Chromosomes of the Sea Lamprey (Petromyzon marinus). Genes. 10(10). doi:10.3390/genes10100832.

Torblaa R, Westman R. 1980. Ecological impacts of lampricide treatments on sea lamprey (Petromyzon marinus) ammocoetes and metamorphosed individuals. Can J Fish Aquat Sci.(37):1835–1850.

Vicoso B. 2019. Molecular and evolutionary dynamics of animal sex-chromosome turnover. Nat Ecol Evol. 3(12):1632–1641. doi:10.1038/s41559-019-1050-8. http://dx.doi.org/10.1038/s41559-019-1050-8.

Vicoso B, Bachtrog D. 2015. Numerous Transitions of Sex Chromosomes in Diptera. PLoS Biol. 13(4):1–22. doi:10.1371/journal.pbio.1002078.

Vicoso B, Emerson JJ, Zektser Y, Mahajan S, Bachtrog D. 2013. Comparative sex chromosome genomics in snakes: differentiation, evolutionary strata, and lack of global dosage compensation. PLoS Biol. 11(8):e1001643. doi:10.1371/journal.pbio.1001643.

Voichek Y, Weigel D. 2020. Identifying genetic variants underlying phenotypic variation in plants without complete genomes. Nat Genet. doi:10.1038/s41588-020-0612-7. http://dx.doi.org/10.1038/s41588-020-0612-7.

Wright AE, Darolti I, Bloch NI, Oostra V, Sandkam B, Buechel SD, Kolm N, Breden F, Vicoso B, Mank JE. 2017. Convergent recombination suppression suggests role of sexual selection in guppy sex chromosome formation. Nat Commun. 8:1–10. doi:10.1038/ncomms14251. http://dx.doi.org/10.1038/ncomms14251.

Wright AE, Dean R, Zimmer F, Mank JE. 2016. How to make a sex chromosome. Nat Commun. 7(May):1–8. doi:10.1038/ncomms12087. http://dx.doi.org/10.1038/ncomms12087.

Yasmin T, Grayson P, Docker MF, Good S V. 2021. The germline-specific region of the sea lamprey genome plays a key role in spermatogenesis. bioRxiv.:2021.09.24.461754. http://biorxiv.org/content/early/2021/09/25/2021.09.24.461754.abstract.

York JR, Lee EM, Mccauley DW. 2019. The lamprey as a model vertebrate in evolutionary developmental biology. In: Lampreys: Biology, Conservation and Control. Springer, Dordecht. p. 481–526.

Zhou X, Stephens M. 2012. Genome-wide efficient mixed-model analysis for association studies. Nat Genet. 44(7):821–824. doi:10.1038/ng.2310.

