## Supplemental Text and Figures for "SexFindR: A computational workflow to identify young and old sex chromosomes"

This PDF file includes:

Supplemental Text (sections 1-3)

Supplementary Table S1

Supplementary Figures S1 to S8

Other supplementary materials for this manuscript include the following:
SupTables_Grayson2022.xlsx, which contains Supplementary Tables S2-S10
(https://github.com/phil-grayson/SexFindR/blob/main/Supplemental_Code/SupTables_Grayson2022.xlsx)

### Supplementary Text

### 1 SexFindR Results

#### 1.1 SexFindR Step 1 - DifCover Results

**Table S1.** DifCover results for 5 species with known sex chromosomes and sea lamprey, which has an unknown sex determination system.

| Species | Sex chromosomes | Total regions analyzed | Total bases analyzed | Male enriched regions | Male enriched bases | Female enriched regions | Female enriched bases | Sex identified |
|---|---|---|---|---|---|---|---|---|
| **Fugu** | Single male-specific SNP (XY) | 1,238 | 3.81e+8 | 211 | 3.99e+6 | 232 | 3.84e+6 | None |
| **Poplar** | 60-100kb Y-like region (XY) | 7,370 | 4.06e+8 | 1,478 | 1.68e+7 | 1,813 | 1.90e+7 | None |
| **Mosquito** | 1.3 Mb M-locus (XY) | 15,035 | 1.26e+9 | 934 | 1.43e+7 | 934 | 1.13e+7 | Male enriched Y |
| **Cannabis** | Slightly expanded Y (XY) | 25,787 | 9.78e+8 | 7,855 | 2.11e+8 | 7,510 | 2.19e+8 | Male enriched Y |
| **Chicken** | Degenerated (ZW) | 2,689 | 1.06e+9 | 387 | 8.55e+7 | 257 | 1.20e+7 | Female enriched W. Male enriched Z. |
| **Lamprey** | Unknown | 10,878 | 1.07e+9 | 1,362 | 1.40e+7 | 1,508 | 1.39e+7 | Unknown |

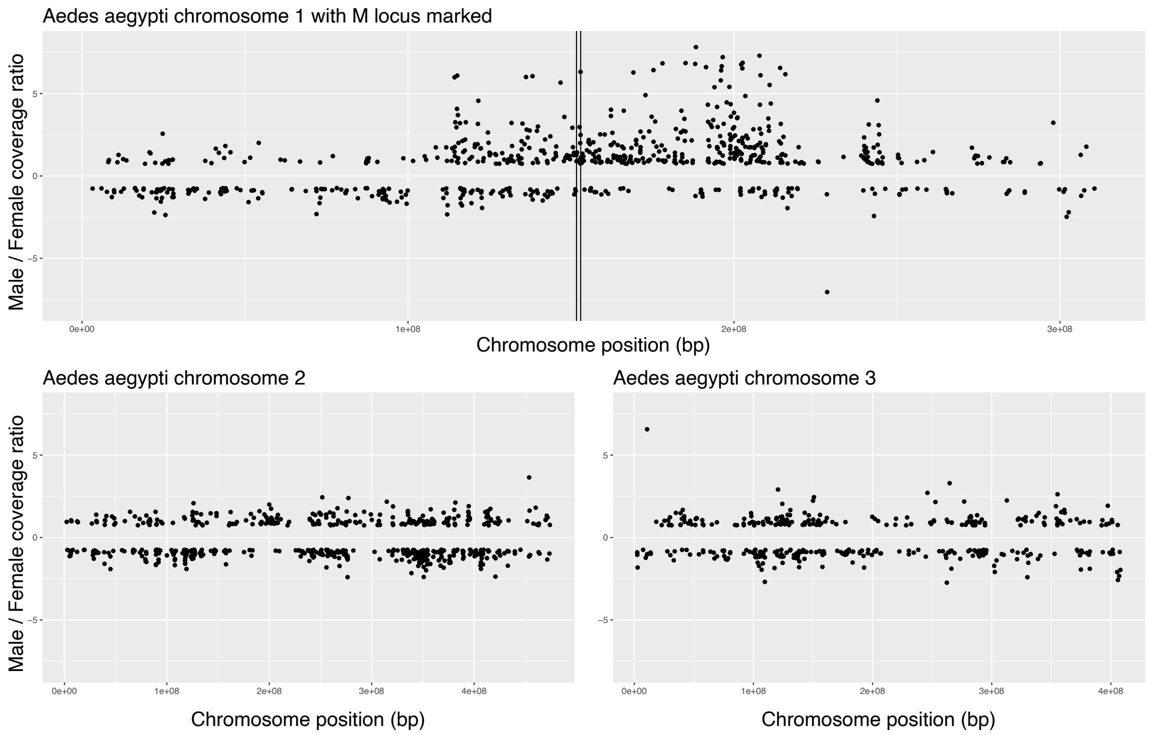


**Fig. S1.** DifCover enrichment scores across the three mosquito chromosomes indicate male-biased coverage across much of the mosquito chromosome 1 surrounding the Y-like M locus (between vertical black lines).

#### 1.2.1 SexFindR Step 2 – Variant-Based Analyses (k-mer)

*Fugu*

As a positive control, the 14 male and 13 female samples from fugu were analyzed using the k-mer pipeline described in the methods. By filtering for the best p-value in the PLINK analysis, we obtained 198 k-mers that appeared to be sex-specific from over 86.5 million initial k-mers that were variable between samples. These were assembled into four small contigs using ABYSS. These sequences all mapped back to NC_042303.1 at the following positions: 9290245-9290305, 12706842-12706902, 12706742-12706844, and 12708070-12708126. The fugu sex determining region (SDR) is found within NC_042303.1:12708070-12708126, and the four other sites found to be fixed between males and females in our dataset fell within these 4 contigs. By viewing this region in IGV, it was found that in our samples, NC_042303.1:9290275 has G/G fixed in all females, while males have either C/G or C/C, and at NC_042303.1:9290314, females are all fixed for A/A, whereas males have either A/T or T/T.  These two sites are within the coding region of fibroblast growth factor receptor substrate 3 (frs3).

We did not find any ABYSS contigs that did not map to the genome, suggesting that sex-specific sequences in fugu are in well-assembled regions. Given the small genome and chromosome level assembly for fugu, this is perhaps not surprising. The pufferfish genome, which is approximately 400 Mb (384 Mb in its current assembly), was first assembled in 2002 and has been greatly improved since (Aparicio et al. 2002; Kai et al. 2011). It contains a similar number of genes to mammals, but much less repetitive sequence – 2.7% interspersed repeats compared to 35-45% in mammals (Aparicio et al. 2002). Given that repetitive sequences are one of the largest challenges for genome assembly, this marked lack of repeats has allowed for a fairly complete genome assembly in fugu.

*Poplar*

kmersGWAS was also run on poplar samples. Due to the disparity in the initial sample number (n=18 males and n=34 females), we subsampled down to (n=18 males and n=19 females) to provide the program with a more balanced distribution of samples. We identified 358,145,003 variable k-mers within these samples. The top p-value out of PLINK was 5.93e-17 and there were 14,530 k-mers that had this significance value. We found that all 14,530 k-mers were male-specific and fully fixed, supporting an XY system. These 14,530 k-mers were assembled with ABYSS into 250 contigs that ranged in size from 28 bp to 175 bp. The majority of these fell within the predicted SDR near the ends of scaffold_19 in the v2.2 poplar genome (fig. S2).

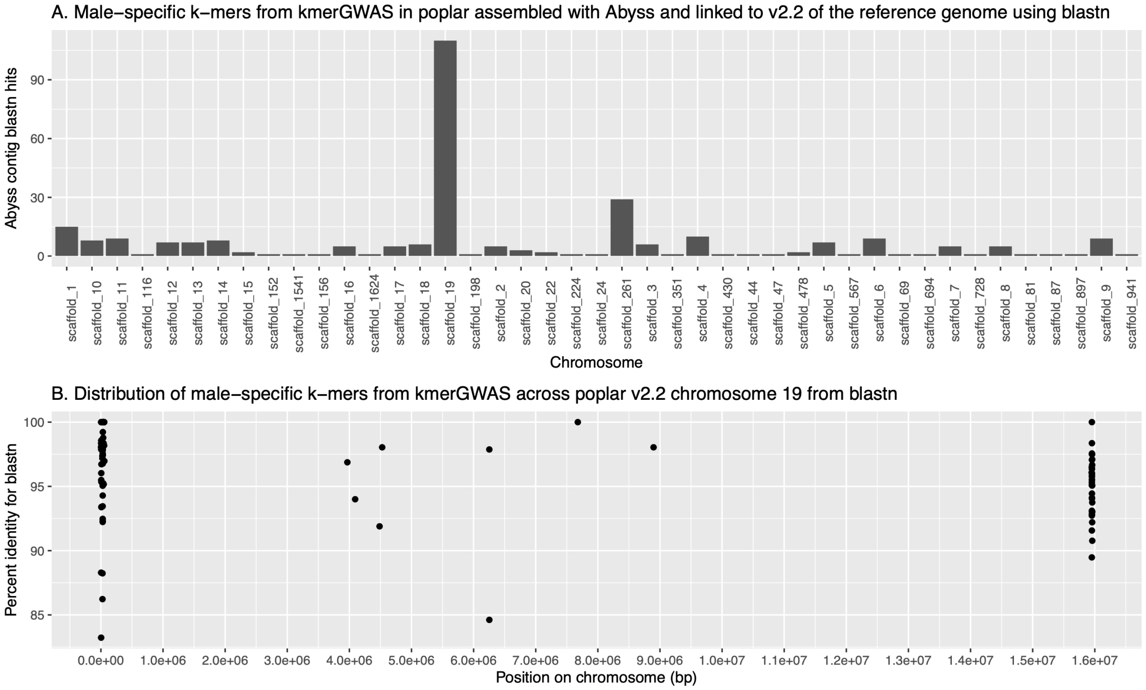


**Fig. S2.** kmersGWAS correctly identified the poplar sex chromosome. (*a*): Male-specific contigs generated from ABYSS map to the v2.2 poplar genome scaffolds using blastn. Scaffold19 (chr19) exhibits the majority of the top hits. (*b*) Distribution of the male-specific blastn hits across chr19. Both the left and right arm of the chromosome have been implicated as SDRs and these contigs primarily overlap those regions.

*Lamprey*

When the k-mer approach was run on the lamprey, over 1 billion variable k-mers were identified within the population and only 56 met our filtering criteria. When assembled with ABYSS, no contigs were created, suggesting that there was very little overlap between the 56 individual k-mers. This lends support to there being no fixed differences between males and females in the sea lamprey.

#### 1.2.2 SexFindR Step 2 – Variant-Based Analyses (GWAS)

*Fugu*

GEMMA (Zhou and Stephens 2012) was run on fugu with 13 females and 14 males, resulting in 1,168,475 analyzed markers. The six top candidate SNPs included the known fugu sex determining region, and they had all a p_lrt value of 8.263923e-204. As a test, a second GEMMA run was conducted where sex was randomly assigned, while maintaining

13 females and 14 males. The same number of markers were analyzed, but the top candidates in the shuffled-sex run had a p_lrt score of 2.738077e-08. These output tables are available on the SexFindR github repository.

*Poplar*

GEMMA analyzed 1,319,754 SNPs across 34 females and 18 males. The top 29 SNPs based on p_lrt, all fell within the SDR in poplar and displayed p_lrt values between 2.306764e-15 and 4.735823e-30.

*Lamprey*

GEMMA was run on the full population data (male n = 126 including both the low and deep sequenced M_8d sample, female n = 140), resulting in analysis of 2,088,765 SNPs. The single top candidate SNP, NC_046112.1:10545622, had a p_lrt score of 2.068937e-08, while remaining SNPs had scores at e-06 or higher (table available on SexFindR github). GEMMA was once again run a second time with randomly shuffled sexes and the top p_lrt score was 1.077499e-06 (table available on SexFindR github). If we filter these true and shuffled datasets for the fugu and lamprey for $p<0.01$, we get 27,849 in sites in the true fugu compared to 14,383 sites in the shuffled fugu, and 27,978 sites in the true lamprey versus 25,552 sites in the shuffled lamprey. Taken together, these data indicate that GEMMA has the power to detect sex-specific changes as small as a single SNP that is fixed between males and females (as was seen in fugu), but that the lamprey data does not offer any immediate promising candidates.

##### 1.2.3 SexFindR Step 2 – Variant-Based Analyses ($F_{ST}$)

*Fugu*

$F_{ST}$ was run on the 13 female and 14 male fugu dataset. A Manhattan plot was generated for the 22 primary chromosomes of fugu (fig. S3), with 5% and 1% outliers marked by horizontal dotted lines and the fugu sex determining region marked by the vertical dotted red line. For fugu, the 1% outliers had $F_{ST}$ values above 0.2475, and 40% of the SNPs above the 1% outlier line fell on the XY chromosome. The fugu SDR was ranked in the top 110 SNPs and the chromosome is an obvious outlier overall.

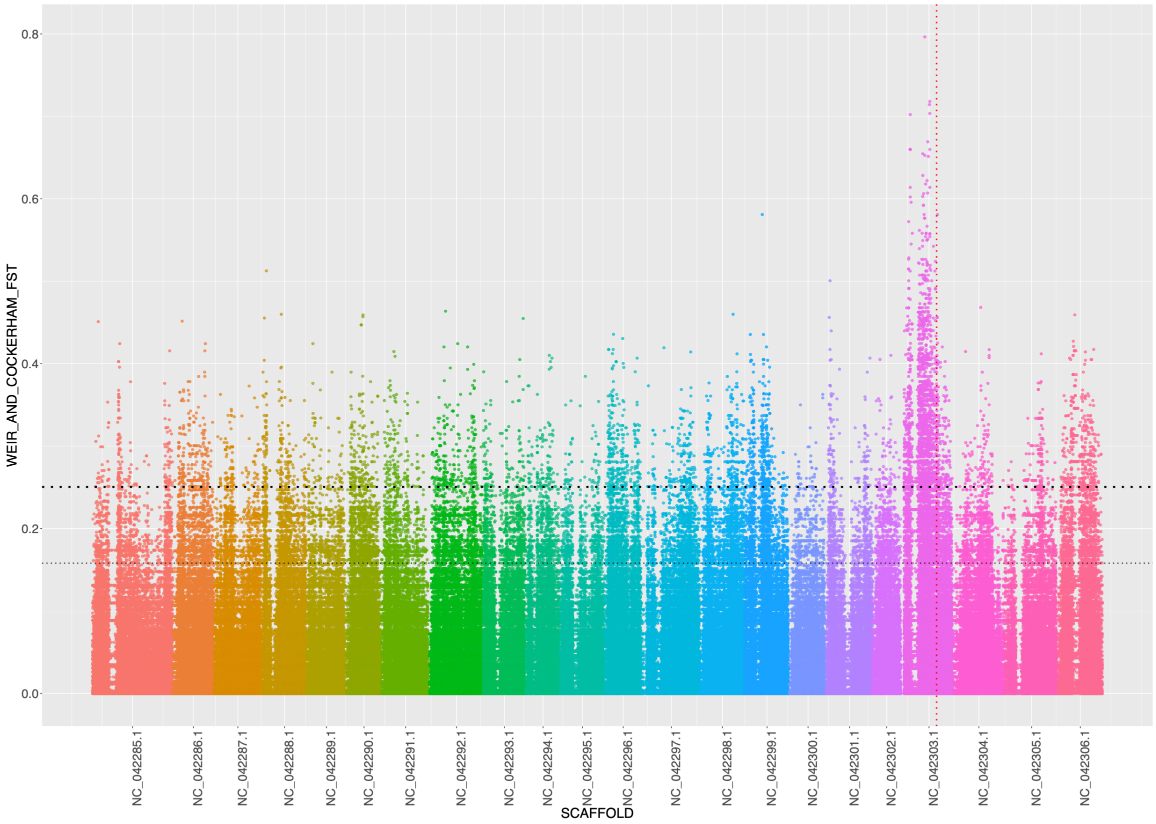


**Fig. S3.** Manhattan plot of $F_{ST}$ values calculated for each SNP on the 22 chromosomes of fugu. The thin horizonal dotted line represents 5% outliers, the thick black dotted line represents 1% outliers, and the vertical red dotted line overlaps the fugu SDR.

*Poplar*

$F_{ST}$ was run on the 34 female and 18 male poplar data set. The same cut-offs were used for a Manhattan plot as used for fugu. Once again, the top candidate sites (n=36) fell within the expected sex-determining region on chromosome 19 (fig. S4).

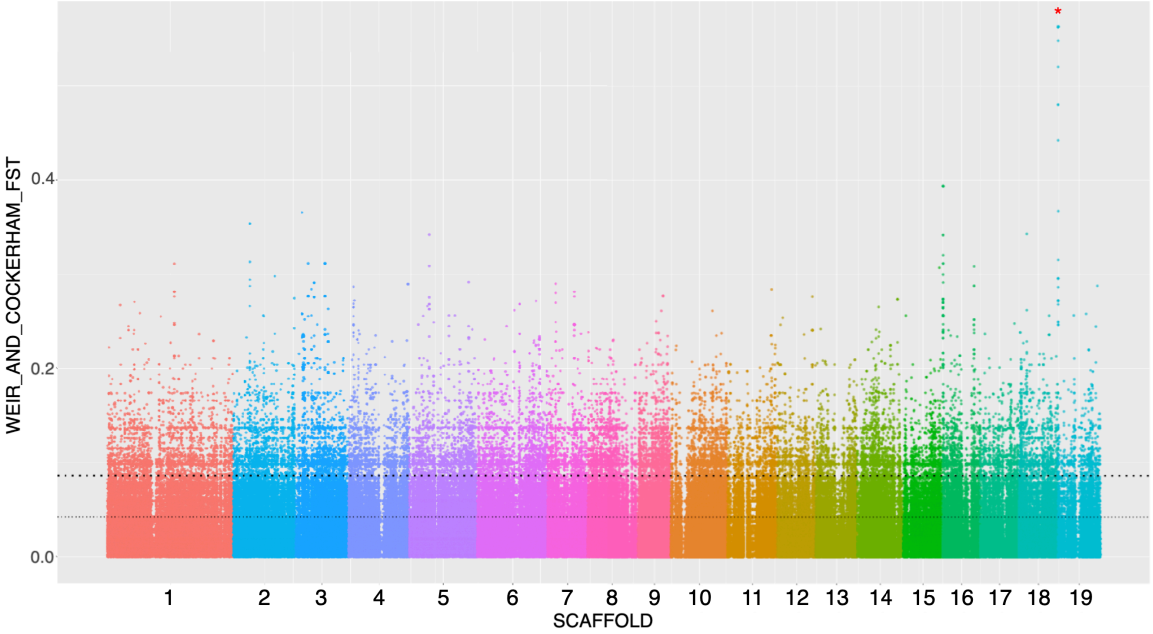


**Fig. S4.** Manhattan plot of $F_{ST}$ values calculated for each SNP on the 19 scaffolds of poplar v2.2. The thin horizonal dotted line represents 5% outliers, the thick black dotted line represents 1% outliers and the highest $F_{ST}$ values are exhibited by sites within the sex determining region, on the left arm of scaffold 19 (region highlighted with red asterisk).

*Lamprey*

A Manhattan plot was generated for 85 assembled scaffolds (fig. S5) and entire genome (fig. S6).

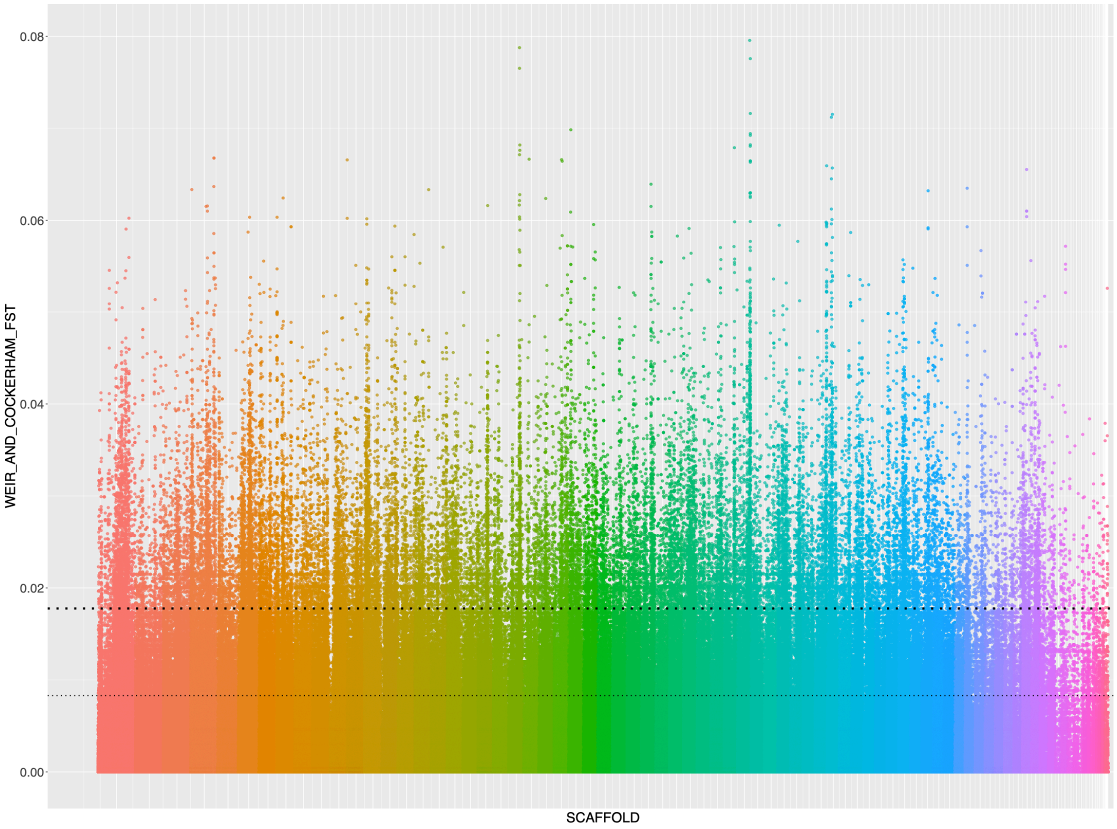


**Fig. S5.** $F_{ST}$ plotted for the 85 assembled chromosomes of the sea lamprey. The thin dotted line designates 5% outliers, whereas the thick dotted line indicates 1% outliers.

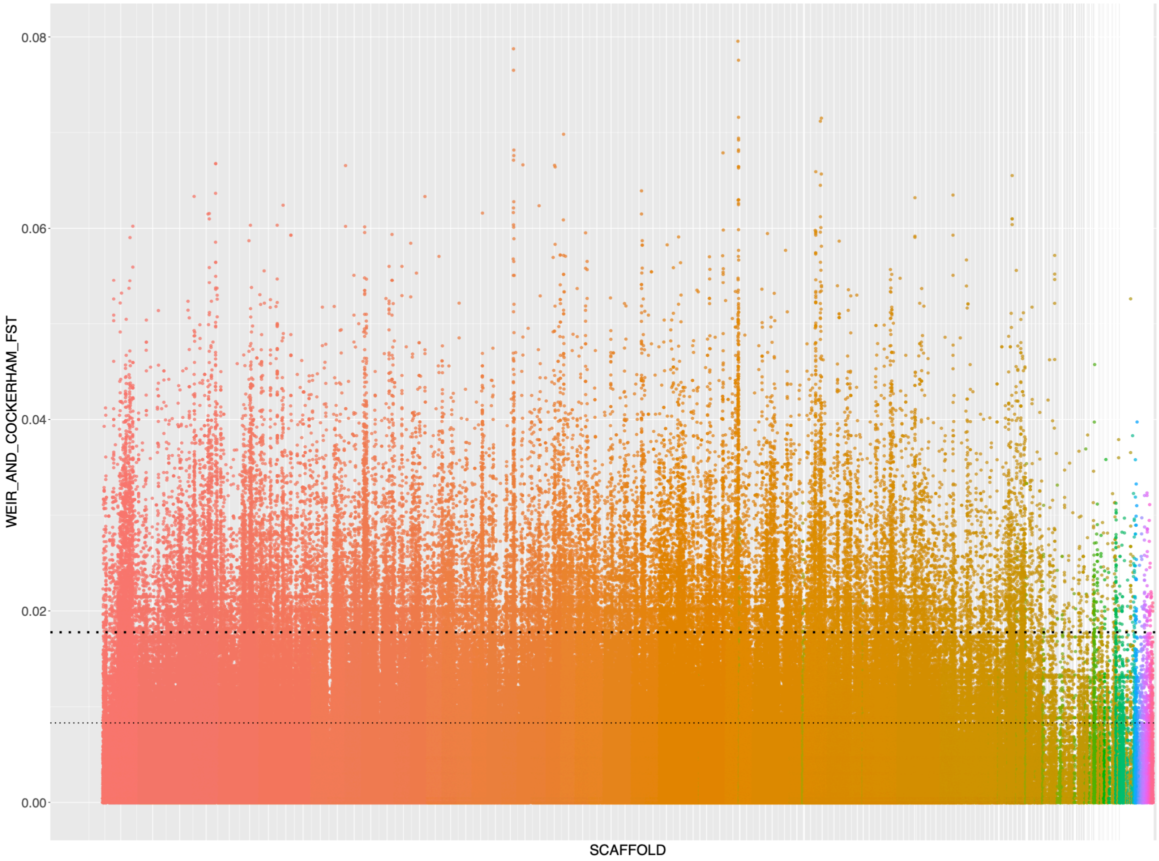


**Fig. S6.** $F_{ST}$ plotted for all chromosomes and scaffolds of the sea lamprey genome. The thin dotted line designates 5% outliers, whereas the thick dotted line indicates 1% outliers.

Since we have simply marked the top 5% and 1% outliers in these Manhattan plots, there is no evidence that sites above these lines have any biological significance in this system. The maximum $F_{ST}$ found in this analysis is just below 0.08, which is very low relative to our other analyses in fugu and poplar. Mean intersexual $F_{ST}$ values were much lower than the positive control species (lamprey mean $F_{ST}$ = 0.0001603402, fugu mean $F_{ST}$ = 0.04949832, poplar mean $F_{ST}$ = 0.02306153). These result suggests that there are not sites that are highly differentiated between the male and female sea lamprey when considered as two separate populations.

#### 1.2.4 SexFindR Step 2 – Variant-Based Analyses (SNP Density)

*Fugu*

SNP density was calculated in 10kb windows for each of the 13 female and 14 male fugu samples. Following the permutations described in the methods section, 37,140 initial windows were analyzed and 4,894 windows (13.18%) reported p-values <= 0.05. The window containing the fugu SDR was among 55 (0.15%) windows with the lowest possible p-value, given 100,000 permutations (0.00000999…). When those windows were ranked by largest male versus female change, the window containing the fugu SDR was in the top 3 windows alongside two other windows on NC_042303.1 that contain sex-specific sequences in our population sample (2,980,000-2,990,000 and 9,260,000-9,270,000).

*Poplar*

SNP density was calculated in 10kb windows for each of the 34 female and 18 male poplar samples. 40,967 windows were run through the 100k permutations, leaving 3,736 (9.1%) that were significant at p <= 0.05 and 344 (0.83%) that displayed the lowest possible p-value in the permutation run. Three of the top four 10kb windows overlapped with the poplar SDR.

*Lamprey*

There were 100,454 initial windows with sex differences in SNP density, with 10,992 (10.94%) windows with p-values <= 0.05, and 627 (0.62%) at p < 0.000999001. The top windows did not yield compelling candidates upon further inspection.

#### 1.3 SexFindR Step 3 – Combined Variant-Based Approach

*Fugu*

Since the three referenced-based population genomics approaches did not perfectly agree for fugu, we tested a combined approach in fugu as a positive control. As described in the main paper, $F_{ST}$, GWAS, and SNP density results were ranked in 10kb windows and their combined signal was used to locate the fugu sex determining region. Rank order was found to be correlated between all three variant-based approaches for fugu. GWAS rank was strongly positively correlated with $F_{ST}$ rank r(*12983*) = 0.6808823, $p < 2.2e\text{-}16$. SNP density rank was weakly positively correlated with both GWAS rank r(3571) = 0.1524189, $p < 2.2e\text{-}16$, and $F_{ST}$ rank r(*2774*) = 0.1545956, $p < 2.592e\text{-}16$. Once all of the ranked lists were assembled in R, a global filter was applied to identify windows that were within the top 100 for GWAS, SNP density and $F_{ST}$. For fugu, from 37,140 initial windows, this filter brought the total number of candidate windows down to five (table S2). One of these five candidate windows contains the fugu SDR, and all five windows were found on the same chromosome, NC_042303.1 (fig. 3).

*Lamprey*

Following the approach outlined above for fugu, 46,366 initial windows produced 10 candidate regions (table S3). Once again, the ranked variant-bases approach produced correlated results, with GWAS and $F_{ST}$ showing the strongest positive correlation r(*30199*) = 0.7912536, $p < 2.2e\text{-}16$. SNP density rank was weakly correlated with both GWAS rank r(*6701*) = 0.1809945, $p < 2.2e\text{-}16$ and $F_{ST}$ rank r(*7404*) = 0.1811085, $p < 2.2e\text{-}16$. In visually assessing these candidate windows, and the SNPs within them, we discovered that the signal appeared to be driven by segregating variation without a strong sex-specific signal. Within these windows, there are no sites across all males or females that were perfectly associated with sex. Although it is possible that a small number of samples could have been sexed or labeled incorrectly in a population data set this large, none of these sites lead us to believe that this would drastically alter these results. This lack of consistency between males and females is also reflected in the relatively low $F_{ST}$ values and the low significance values associated with individual SNPs in the GWAS. However, these candidate regions could possess some biological signal related to sex determination or sexual antagonism. It is also possible that all of these regions are simply artifacts of the filtering strategy given that we have chosen an arbitrary cut-off of the top 5% for the $F_{ST}$ and GWAS results, and these analyses seem to produce correlated results. None of these regions overlapped with well-documented sex determining genes from other species (table S9).

### 2 Supplementary Lamprey Analyses

#### 2.1 DifCover 2-by-2 analysis

We identified an outlier scaffold in the Step 1 coverage analyses in lamprey (fig. 2), which could indicate sex-specific sequence. To assess the possibility of large region of significant sequence divergence between males and females on a lamprey scaffold, we conducted follow-up screenings.

First, we analyzed multiple DifCover runs together for original Huron lamprey samples and validated this method with chicken. Bedtools makewindows was used to generate 10kb windows across the genome and bedtools annotate was used to identify genomic regions that were consistently male- or female-biased across 4 experimental DifCover runs (F1/M1, F1/M2, F2/M1, and F2/M2) and not enriched in either control DifCover run (F1/F2 and M1/M2). In chicken, 106,788 10kb windows were analyzed. This resulted in 7,396 male-enriched windows across 6 scaffolds with 7,390 of them falling on the Z, and 681 female-enriched windows across 30 scaffolds, with 626 of them falling on the W. In lamprey, 109,630 windows were analyzed. This resulted in 186 male-enriched windows across 64 scaffolds, with NC_046134.1 having the most (n=19 windows), and 77 female-enriched windows across 39 scaffolds, with NC_046128.1 having the most (n=5 windows). The major outlier scaffold in Figure 2*f* (NC_04153.1) was only present in the one pairing of samples F2 and M6, and was eliminated as a possible candidate region. Finally, when the low coverage original Huron lamprey samples (n = 8 males, n=8 females) were analyzed with DifCover and their significant regions of differential coverage were used to filter the high coverage dataset, the number of male-enriched windows decreased to 30 windows across 7 scaffolds, with all 19 windows for NC_046134.1 remaining significant, and only 6 windows across 5 scaffolds for female-enriched regions.

While assessing the only major candidate from the extended lamprey DifCover analyses (19 windows of significantly deeper male sequencing on NC_046134.1), we examined this region across all sequenced populations and discovered what appears to be a segregating deletion that results in a false positive signal (fig. S7). Across all populations analyzed, there is a noticeable drop in coverage for both males and females within the highlighted region, indicating that the deletion is very common (fig. S7*a*). Across all populations, the homozygous reference (no deletion) genotype is the rarest, which allows a small number of individuals (males or females) in the population to drive the false positive signal.

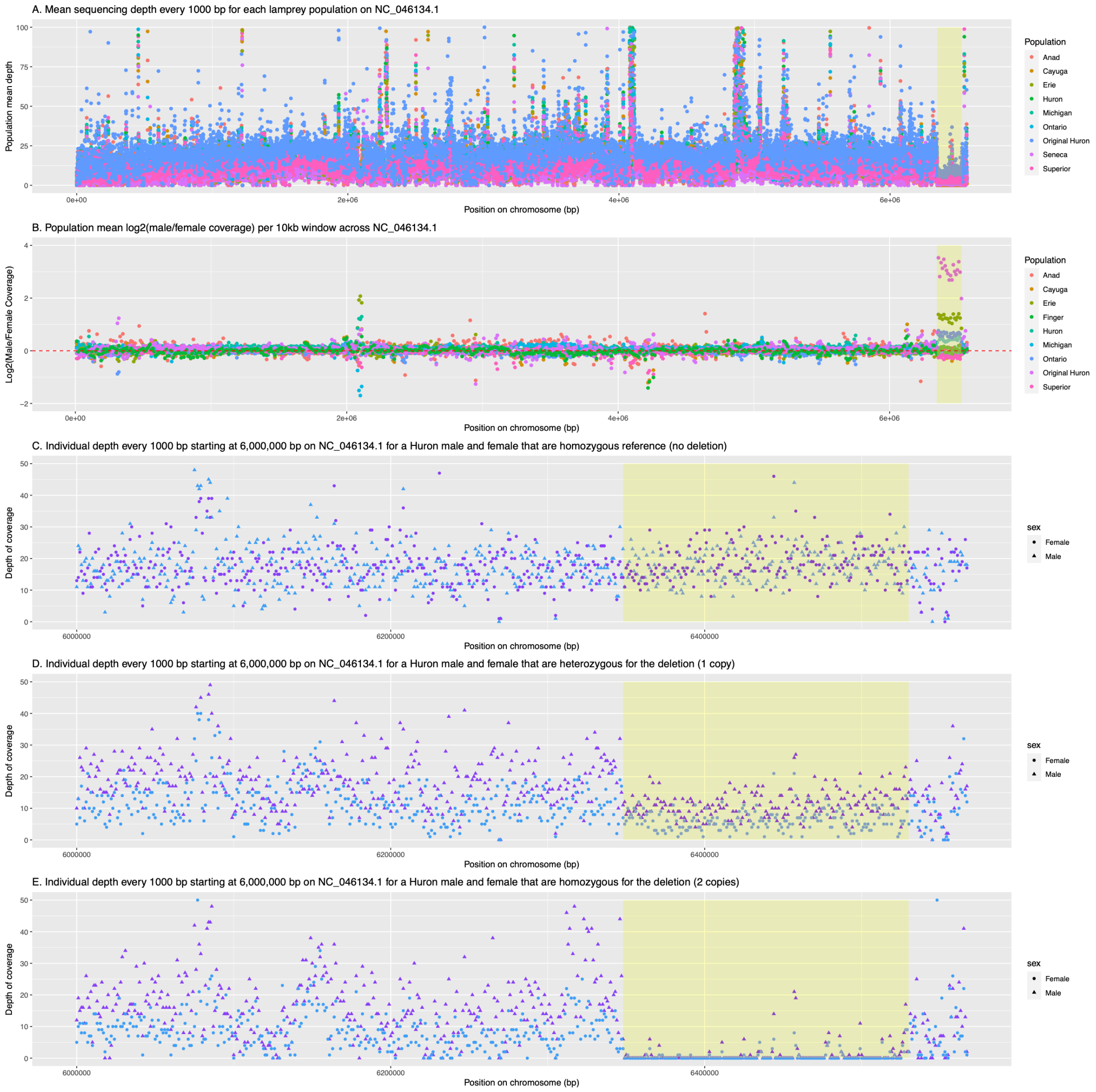


**Fig. S7.** Evidence for a large segregating deletion from ~6,348,000-6,530,000 bp on NC_046134.1 which underpins the false positive signal in coverage-based analyses. A: Across all populations analyzed there is a notable drop in average sequencing coverage throughout this region (max coverage limited to 100x for plot). B: Slight differences in the number of males or females with the segregating deletion can provide false positive signal in coverage-based analyses across different populations. C-E: One male and one female that display each possible genotype were selected from Lake Huron drainage populations to demonstrate that the deletion is not sex-specific (max coverage was limited to 50x for these plots).

#### 2.2 Testing for population-specific sex-linked regions in the lamprey

Given the large number of populations examined together in the lamprey analyses, we wanted to test whether there could be sex-specific SNPs that were only population-specific. An in-house python script (<species>_vcf_search.py) was written to search the VCF file for sites that contained sex-specific differences within a single population. To test this method, it was first run on fugu (fugu_vcf_search.py), where it identified 6 sites that have perfect correlation between males and females in the VCF file, including the fugu SDR (table S10).

For the lamprey, this script (lamprey_vcf_search.py) was run on the following populations: Cayuga Lake (male n= 11, female n= 9), Lake Erie (male n= 16, female n= 14), Holyoke, MA, Holyoke, MA anadromous fish (male n= 7, female n= 5), Lake Huron (male n=39, female n=41), Lake Michigan (male n= 13, female n= 17), Lake Ontario (male n= 16, female n= 22), original Huron samples (male n= 9, female n= 11), and Lake Superior (male n= 16, female n= 31). From all these populations, only the anadromous fish from Holyoke had any sites that were found to be sex-specific, but with data from only seven males and five females, it is unlikely that these are truly fixed in the population. To test the significance of these potential sites, GEMMA was run on only the anadromous population. This resulted in 1,783,178 SNPs being analyzed with the top hits displaying p_lrt values of 3.065630e-95. When sex was randomly assigned for a second anadromous GWAS while maintaining the number of samples that were assigned to the different sexes (male n= 7, female n= 5), top hits obtained the same significance value (3.065630e-95), and the distribution of sites within the different p_lrt bins was nearly identical to the true run, suggesting that this signal is an artifact of low sample number within the population and has no biological relevance (fig. S8).

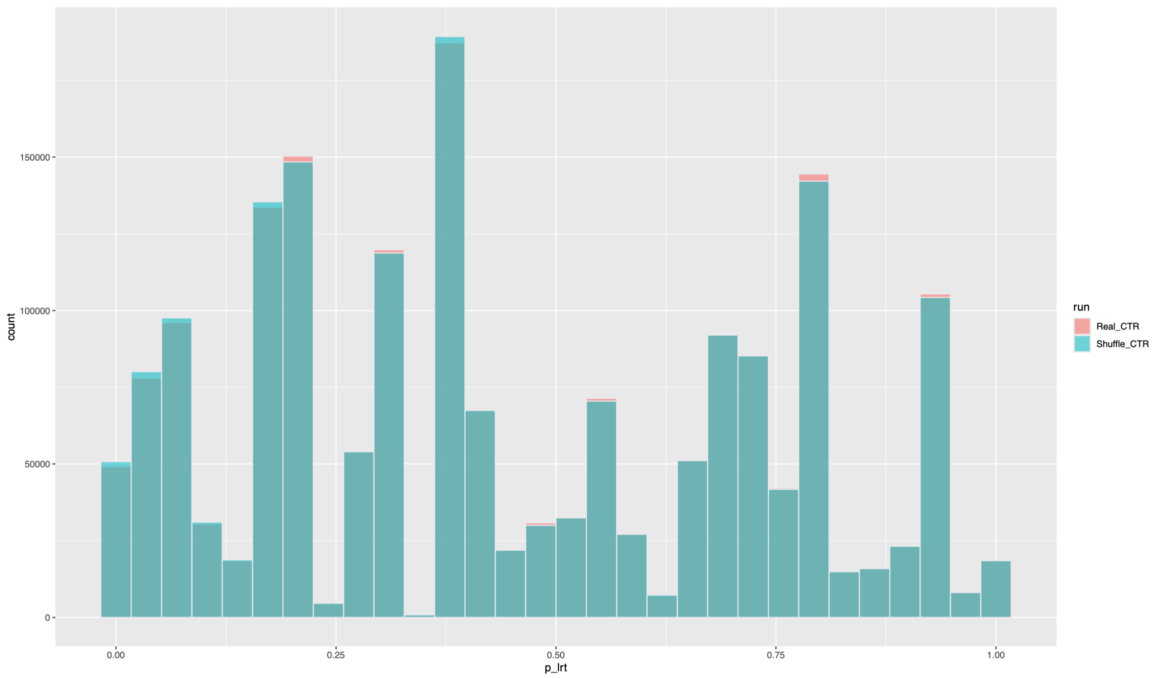


**Fig. S8.** Stacked histogram of p_lrt values in the true anadromous samples (Real_CTR) and the sex-shuffled anadromous samples (Shuffle_CTR) following a GEMMA GWAS on each group. This nearly identical distribution of p_lrt values indicates that the GWAS has low power with low sample number and the fixed differences identified at the population level in the true data with male n=7 and female n=5 are not significantly different from a null expectation with low sample numbers.

### 3 Supplementary References

Aparicio S, Chapman J, Stupka E, et al (2002) Whole-genome shotgun assembly and analysis of the genome of *Fugu rubripes*. Science (80- ) 297:1301–1310. https://doi.org/10.1126/science.1072104

Kai W, Kikuchi K, Tohari S, et al (2011) Integration of the genetic map and genome assembly of fugu facilitates insights into distinct features of genome evolution in teleosts and mammals. Genome Biol Evol 3:424–442. https://doi.org/10.1093/gbe/evr041

Zhou X, Stephens M (2012) Genome-wide efficient mixed-model analysis for association studies. Nat Genet 44:821–824. https://doi.org/10.1038/ng.2310
